# AI-based Predictive Signaling Pathway Profiling in Cardiac Fibrosis Suggests a Novel Combinatorial Treatment Strategy

**DOI:** 10.64898/2025.12.04.692460

**Authors:** Bo Yang, Jie Chen, Qiongjie Mi, Jiuzhou Huo, Ningxin Gao, Xinhua Lin, Jeffery D. Molkentin, Qinghang Meng

## Abstract

**Background:** Cardiovascular disease (CVD) remains the leading cause of global mortality, with myocardial fibrosis characterized by excessive extracellular matrix (ECM) deposition representing a common endpoint associated with progressive cardiac dysfunction. While studies in animal models of heart disease suggest that preventing or reducing fibrosis can antagonize negative ventricular remodeling, current therapeutic strategies remain clinically limited and ineffective, in part due to an incomplete understanding of the multifactorial nature of the fibrotic process.

**Methods and Results:** We employed artificial intelligence (AI) trained from 6,528 cardiac fibrosis publications over the past decade, which suggested 10 nodal signaling pathways underlying the fibrotic process. Single-cell RNA sequencing was used to quantify pathway activities across mouse models and clinical samples encompassing acute and chronic cardiac injury. Dynamic enrichment analysis revealed 2 critical pathways involved in acute myocardial infarction (MI) injury driven temporally-regulated fibrosis whereby Janus kinase - Signal transducer and activator of transcription (JAK-STAT) signaling peaked early (day 7) while transforming growth factor-β (TGF-β) pathway activity peaked mid-phase (day 14). With more chronically-driven fibrosis, JAK-STAT signaling again emerged, which this time was persistently active during chronic transverse aortic constriction (TAC) injury. Experimentally, single-cell analysis identified a pathogenic myofibroblast subpopulation characterized by high JAK-STAT signaling and ECM secretion capacity. Fibroblast-specific *Jak1/2* gene-deleted mice significantly reduced TAC-induced fibrosis by blocking myofibroblast formation from this subpopulation, although these same mice failed to show attenuated fibrosis following MI injury, suggesting a pathway that would be ideal to therapeutically target for chronic fibrosis without affecting necessary acute scar formation. Indeed, based on the predictive AI driven algorithm, a temporal inhibitory strategy was generated for JAK-STAT and TGF-β that permitted compensatory scar formation while more effectively preventing progressive fibrosis and worsened cardiac function versus either singular pathway or chronic antagonism.

**Conclusion:** Here we employed the wealth of past signaling pathway data underlying cardiac fibrosis to train an AI model, which suggested an optimized approach of inhibiting 2 nodal signaling pathways but with differential timing as a more effective therapeutic strategy that reduces pathologic cardiac fibrosis without negatively impacting compensatory fibrotic activity with acute MI injury. The therapeutic approach involves altering the timing of JAK-STAT signaling blockade with ruxolitinib (RUX) in combination with delayed and temporary TGF-β signaling inhibition post MI injury with pirfenidone (PFD).

## Introduction

Cardiac fibrosis represents a pathological hallmark characterized by excessive extracellular matrix (ECM) deposition within the myocardial interstitium, leading to increased ventricular stiffness and progressive deterioration of cardiac function^1–5^. This process constitutes a common endpoint across diverse cardiovascular diseases (CVDs), including myocardial infarction (MI), cardiomyopathy, and heart failure^6–10^. Despite decades of intensive research, clinical interventions targeting cardiac fibrosis have yielded disappointing results^11,12^. This therapeutic failure stems primarily from the dynamic and context-dependent nature of key molecular targets during fibrotic progression. Myofibroblasts, the principal effector cells driving scar formation^13^, exhibit a narrow proliferative window of approximately one week post-MI, with their population declining by two-thirds from peak levels during later stages of infarction^14^. Similarly, while the transforming growth factor-β (TGF-β)/SMAD family member 3 (SMAD3) pathway represents a well-validated therapeutic target, its clinical translation remains elusive^11,15–18^. Preclinical studies have demonstrated that premature or prolonged TGF-β inhibition increases mortality in MI models, emphasizing the critical importance of intervention timing^19^. These observations highlight that even thoroughly characterized targets require precise temporal modulation to achieve therapeutic efficacy.

The temporal complexity of cardiac fibrosis extends beyond individual pathways to encompass pathway interactions and context-dependent functions. Identical signaling cascades may exhibit opposing roles depending on disease stage, cellular context, and pathological environment. Furthermore, the sequential activation of multiple pathways during fibrotic progression suggests that combination therapies targeting complementary mechanisms may prove more effective than single-pathway approaches. Indeed, past data interrogation and predictive modeling has suggested critical nodal regulators of cardiac fibroblast activation and disease-based fibrosis, which was further refined to suggest combinatorial therapeutic strategies^20–26^, although temporal modeling of maximal pathway activity was not fully developed.

To address these challenges, we developed an integrated research framework examining three critical dimensions of cardiac fibrosis therapeutics which are target heterogeneity across different pathological contexts, dynamic regulatory windows governing pathway activation, and opportunities for temporally coordinated combination therapy. Our approach encompassed multi-dimensional data integration through bibliometric analysis of 6,528 publications and reanalysis of single-cell transcriptomic datasets from independent studies^14,27,28^. We constructed spatiotemporal atlases mapping dynamic enrichment patterns of 10 signaling pathways across mouse models of acute and chronic injury, as well as three clinical sample types.

Our findings demonstrate that Janus kinase - Signal transducer and activator of transcription (JAK-STAT) inhibition ameliorates chronic cardiac fibrosis but exhibits no efficacy in acute MI when specifically targeted in myofibroblasts, illustrating pathway-specific therapeutic windows. Based on these temporal activity patterns, we developed a combinatorial strategy employing early-phase JAK-STAT inhibition followed by mid-phase TGF-β inhibition. Preclinical validation confirmed that this temporally coordinated approach significantly outperformed single-pathway interventions in reducing scar formation, attenuating fibrosis, and preserving cardiac function.

Our study establishes the fundamental importance of temporal dynamics in cardiac fibrosis therapeutics. Our results demonstrate that the failure of traditional single-target approaches often results from misaligned intervention timing rather than inappropriate target selection. By precisely coordinating therapeutic interventions with pathway activity peaks, temporally optimized combination therapy offers a promising strategy to overcome current limitations in antifibrotic treatment.

## Materials and Methods

### Data Availability

The data, analytic methods, and study are available from the corresponding author on reasonable request.

### Mice

All animal procedures were approved by the Institutional Animal Care and Use Committee (IACUC) of the Greater Bay Area Institute of Precision Medicine (Guangzhou). Age-matched male and female C57BL/6 mice were used and randomized across experimental groups to ensure balanced sex distribution. To generate mice with fibroblast-and myofibroblast-specific deletion of *Jak1* and *Jak2*, we crossed *Jak1/2^flox/flox^*mice^29^ with *Rosa26^NG/wt^* reporter mice^13^ and either *Periostn^MCM/+^* mice^13^ or *Pdgfrα^creERT^* mice (Jaxson Laboratory, stock #018280). All breeding pairs were littermate-controlled to minimize genetic variability. For Cre-mediated gene deletion, tamoxifen (75 mg/kg/day) was administered intraperitoneally for 14 consecutive days to simultaneously delete *Jak1* and/or *Jak2* while activating *Rosa26^NG/wt^* reporter expression. PCR-based genotyping was performed to confirm the presence of floxed alleles, Cre recombinase, and the reporter construct in all experimental animals.

### Drug Administration

Ruxolitinib (RUX; 50 mg/kg, catalog #V0076, Invivochem) was dissolved in vehicle solution (2% DMSO, 30% PEG300, 68% ddH₂O) and administered via oral gavage twice daily with ≥8-hour intervals. Treatment began 24 hours prior to surgery and continued until endpoint assessment. Control mice received vehicle alone using the same administration schedule. Three therapeutic regimens were evaluated. (1) RUX monotherapy: mice received RUX (50 mg/kg, twice daily) from 1 day pre-surgery through postoperative day 8. (2) Pirfenidone (PFD) monotherapy: mice were administered PFD (150 mg/kg in 50% PEG300 and 50% saline; catalog #HY-B0673, MedChemExpress) once daily from postoperative day 6 through study endpoint. (3) RUX+PFD combination therapy: this sequential treatment protocol consisted of RUX monotherapy (1 day pre-surgery through postoperative day 8), followed by RUX+PFD co-administration (postoperative days 6-8), and finally PFD monotherapy alone (postoperative day 9 through endpoint).

### Transverse Aortic Constriction (TAC) Surgery

All surgical procedures were performed under sterile conditions. Mice were anesthetized with isoflurane (1.5-2% in 100% O₂) and intubated to maintain controlled ventilation. The surgical site was shaved and disinfected with povidone-iodine, followed by sterile draping. A midline cervical incision was made, and the thoracic cavity was accessed through partial clavicular excision and separation of the second and third ribs under stereomicroscopic guidance. The thyroid gland and perivascular fat were carefully retracted to expose the transverse aortic arch. TAC was performed by placing a 6-0 polypropylene suture around the aorta between the brachiocephalic trunk and left common carotid artery. A blunt 27-gauge needle was used as a caliber reference during ligation to ensure standardized constriction. The needle was removed immediately after suture placement, creating reproducible pressure overload. The thoracic wall was closed in layers using 6-0 nylon sutures (sternum first, followed by skin closure). Mice were maintained at 37°C and monitored until full recovery of spontaneous ventilation was achieved. Sham-operated controls underwent identical surgical procedures without aortic ligation. All mice received sustained-release buprenorphine (0.2 mg/kg, subcutaneous) for postoperative analgesia.

### Myocardial Infarction (MI) Surgery

All surgical procedures were performed under sterile conditions. Mice were anesthetized with 1% sodium pentobarbital (0.1 mL per 25 g body weight, intraperitoneal injection) and intubated, then positioned in right lateral decubitus. The anterior chest and left axilla were shaved and disinfected with povidone-iodine, followed by sterile draping. A transverse incision (∼2 cm) was made in the left axilla with blunt dissection of the pectoral muscles to expose the second and third intercostal spaces. The thoracic cavity was accessed via resection of intercostal muscles and ribs using micro-scissors. The pericardium was carefully opened to visualize the left anterior descending (LAD) coronary artery. MI was induced by permanent ligation of the mid-LAD using 7-0 polypropylene suture. Successful ligation was confirmed by immediate myocardial pallor and anterior left ventricular wall dyskinesia. The thoracic cavity was closed in layers using 6-0 nylon sutures followed by continuous skin closure. Mice were maintained at 37°C and monitored until recovery of spontaneous ventilation. Sham-operated controls underwent identical surgical procedures without LAD ligation. All mice received sustained-release buprenorphine (0.2 mg/kg, subcutaneous) for postoperative analgesia.

### Echocardiography

Transthoracic echocardiography was performed preoperatively (baseline) and postoperatively on mice subjected to TAC or MI using a high-resolution ultrasound system (VINNO 6, VINNO Technology, China) equipped with a 35-MHz linear-array transducer. To ensure physiological stability during imaging, mice were anesthetized with 1.5-2.0% isoflurane in 100% oxygen delivered via a nose cone, maintaining core body temperature at 37±0.5°C using a heated platform. Heart rate was monitored continuously and maintained between 400-600 beats per minute (bpm), consistent with murine physiological norms under light anesthesia. Mice were positioned in left lateral decubitus, and images were acquired in the parasternal long-axis (PLAX) view to optimize alignment of the left ventricle (LV). M-mode recordings were obtained at the level of the papillary muscles to measure: Left ventricular internal diameter (LVID), left ventricular posterior wall thickness (LVPW) and interventricular septal thickness (IVS). Measurements were performed at end-diastole and end-systole, defined respectively as the maximal and minimal LV chamber dimensions. Three consecutive cardiac cycles were analyzed per animal, and values were averaged to minimize beat-to-beat variability.

### Flow Cytometry

Single-cell suspensions were prepared from freshly isolated mouse ventricles (both male and female) using an enzymatic dissociation protocol^13^. Briefly, ventricular tissues were minced into 1-2 mm³ fragments and subjected to three sequential digestions at 37°C in an enzymatic solution containing: Collagenase type II (2 mg/mL, C2-28, Sigma), Dispase II (1.2 U/mL, D4693, Sigma), 0.9 mM CaCl₂ and DMEM supplemented with 2% FBS and antibiotics. Digestion was performed with progressive pipette size reduction (10 mL, 5 mL and 1 mL serological pipettes) between 20-minute incubation intervals. Cell suspensions were filtered through 40 μm mesh, erythrocytes were lysed (C3702, Beyotime), and cells were resuspended in sorting buffer (HBSS with 2% bovine growth serum and 2 mM EDTA). For immunophenotyping, cells were stained with fluorescent-conjugated antibodies against: CD31 (102408, BioLegend), CD45 (103116, BioLegend) and MEF-SK4 (130-120-802, Miltenyi). All samples were analyzed on a BD FACSAria III flow cytometer (BD Biosciences), with centrifugation steps performed at 400 ×g (10 min, 4°C) using a swinging bucket rotor without brake application.

### Single-cell RNA sequencing and analysis

Single-cell transcriptomes were generated using the 10x Genomics Chromium Single Cell 3’ Reagent Kit v3. Briefly, viable cells (≥85% viability) were resuspended at 700-1200 cells/μL in 0.04% BSA-DPBS and partitioned into Gel Bead-in-Emulsions (GEMs) via microfluidics, achieving >90% single-cell capture efficiency. Reverse transcription within GEMs produced barcoded cDNA, which was subsequently amplified and fragmented for library construction. Sequencing was performed on an Illumina NovaSeq platform (150 bp paired-end reads). Raw data were processed using Cell Ranger (v6.1.2) for demultiplexing, alignment (GRCh38 reference genome), and UMI counting. Downstream analyses employed Seurat (v4.0): cells expressing <200 genes or with>10% mitochondrial reads were excluded, followed by normalization (SCTransform).

### Bulk RNA sequencing

Approximately 1 × 10⁶ sorted cells were pelleted by centrifugation, lysed in 1 mL TRIzol reagent (15596018; ThermoFisher), and gently pipetted to minimize mRNA degradation. After 5 min of incubation at RT (allowing complete dissociation of nucleoprotein complexes), 0.2 mL chloroform was added, followed by vigorous shaking for 15s and a 2-3 min incubation. The mixture was centrifuged at 12,000 × g for 15 min at 4°C, and the upper aqueous phase containing RNA was transferred to a fresh tube. RNA was precipitated by adding an equal volume of isopropanol (I112011; Aladdin), incubating for 10 min at RT, and centrifuging at 12,000 × g for 10 min at 4°C. The pellet was washed with 1 mL of 75% ethanol (E809056; Macklin), centrifuged at 7,500 × g for 5 min at 4°C, and air-dried for 5-10 min (avoiding complete desiccation). Purified RNA was resuspended in 30-50 μL DEPC-treated water. RNA concentration and integrity were assessed using the Agilent TapeStation system, with RNA Integrity Numbers (RIN) ranging from 8.0 to 10.0. Poly(A)+ RNA was enriched via two rounds of oligo (dT) selection and reverse-transcribed into cDNA. Sequencing libraries were prepared using the VAHTS Universal V8 RNA-seq Library Prep Kit for Illumina (NR605-02; Vazyme) and quantified by qPCR. Paired-end sequencing (2 × 150 bp) was performed on the Illumina HiSeq 2500 platform, achieving an average depth of >30 million reads per sample. Raw reads were aligned to the mm10 reference genome using the STAR aligner (v2.7.10a) within the bcbio-nextgen RNA-seq pipeline (v1.2.4). Differential gene expression analysis was conducted using DESeq2 (v1.30.1) in R (v4.0.3). Single-cell RNA and bulk RNA sequencing were performed by BerryGenomics Co., Ltd.

### Histology

Following euthanasia, mouse hearts were rapidly excised and immediately immersed in 4% paraformaldehyde (PFA) solution containing 0.2mM potassium chloride (KCl) for 24 h at 4°C to ensure optimal fixation while preserving cardiac architecture. Fixed tissues were dehydrated through a graded ethanol series (30%, 50%, 70%, 85%, 95%, and 100%), cleared in xylene, and embedded in paraffin for sectioning. Serial sections (5 μm thickness) were obtained using a microtome and mounted on positively charged glass slides for subsequent staining. Heart sections were stained using a Masson’s Trichrome kit (G1006; Servicebio) following the manufacturer’s protocol. Collagen fibers were visually identified as blue-stained regions, while cardiomyocytes and cytoplasm appeared red. Quantification of fibrosis was performed using ImageJ (NIH, v1.54) with color thresholding to determine the percentage of fibrotic area (blue) relative to total tissue area per sample. To evaluate cardiomyocyte hypertrophy, sections were incubated with fluorescein-conjugated wheat germ agglutinin (W849; Invitrogen) for 1 h at room temperature, delineating myocyte membranes. Nuclei were counterstained with DAPI (40727ES10; Yeasen). Cardiomyocyte surface area (CSA) was quantified in transversely cut cardiomyocytes using ImageJ. A minimum of 100 myocytes per mouse were measured, and the mean CSA was calculated. All analyses were performed in a blinded manner by two independent investigators. Data are presented as mean ± SD, with sample sizes (n) indicated in figure legends.

## Statistical Analysis

Statistical analyses were performed using GraphPad Prism 9.5 (GraphPad Software). Data are presented as mean ± SD. One-way ANOVA with Tukey’s post hoc test was used for multiple group comparisons. Survival analysis was conducted using Kaplan-Meier curves with log-rank (Mantel-Cox) tests. All tests were two-tailed, with *p* < 0.05 considered statistically significant.

## Results

### Temporal dynamics of core fibrosis-related signaling pathways across cardiac disease progression

To systematically identify the core regulatory mechanisms underlying cardiac fibrosis, we performed a comprehensive bibliometric analysis of fibrosis-related signaling pathways. We retrieved publications from the PubMed database spanning July 2015 to July 2025 using the R package *easyPubMed*, identifying 6,528 articles containing “cardiac fibrosis” in either the title or abstract. Through natural language processing, we quantified the occurrence frequency of 16 KEGG pathway terms and ranked them by weighted frequency (Figure 1A and Table S1). Our analysis revealed that the TGF-β signaling pathway (KEGG ID: hsa04350) and ECM-receptor pathway (KEGG ID: hsa04512) were the most extensively investigated, with occurrence frequencies of 2,132 (32.7%) and 1,041 (15.9%), respectively. This distribution is consistent with established research paradigms that the TGF-β pathway functions as a central regulator of fibrosis by promoting cardiac fibroblast activation through SMAD2/3 phosphorylation^15,30,31^, while the ECM-receptor pathway directly reflects the terminal fibrotic phenotype, with accumulation of its core components (COL1A1, COL3A1, FN1, POSTN) serving as the gold standard for pathological diagnosis^32,33^. Notably, type I/III collagen (COL1A1/COL3A1) is predominantly secreted by activated myofibroblasts and constitutes the structural foundation of cardiac fibrosis^34^. Additional highly frequent pathways included Apoptosis, MAPK, WNT, FGFR, JAK-STAT, Chemokine, Cell Cycle, and P53 signaling pathways. To specifically examine the role of fibroblasts in cardiac fibrosis, we performed a secondary analysis incorporating the keyword “fibroblast” [query: “cardiac fibrosis” (Title/Abstract) and “fibroblast” (Title/Abstract)], which yielded 1,159 articles. The top five pathways remained unchanged (Figure S1A), further confirming the central role of fibroblasts in cardiac fibrosis pathogenesis.

**Figure 1.**
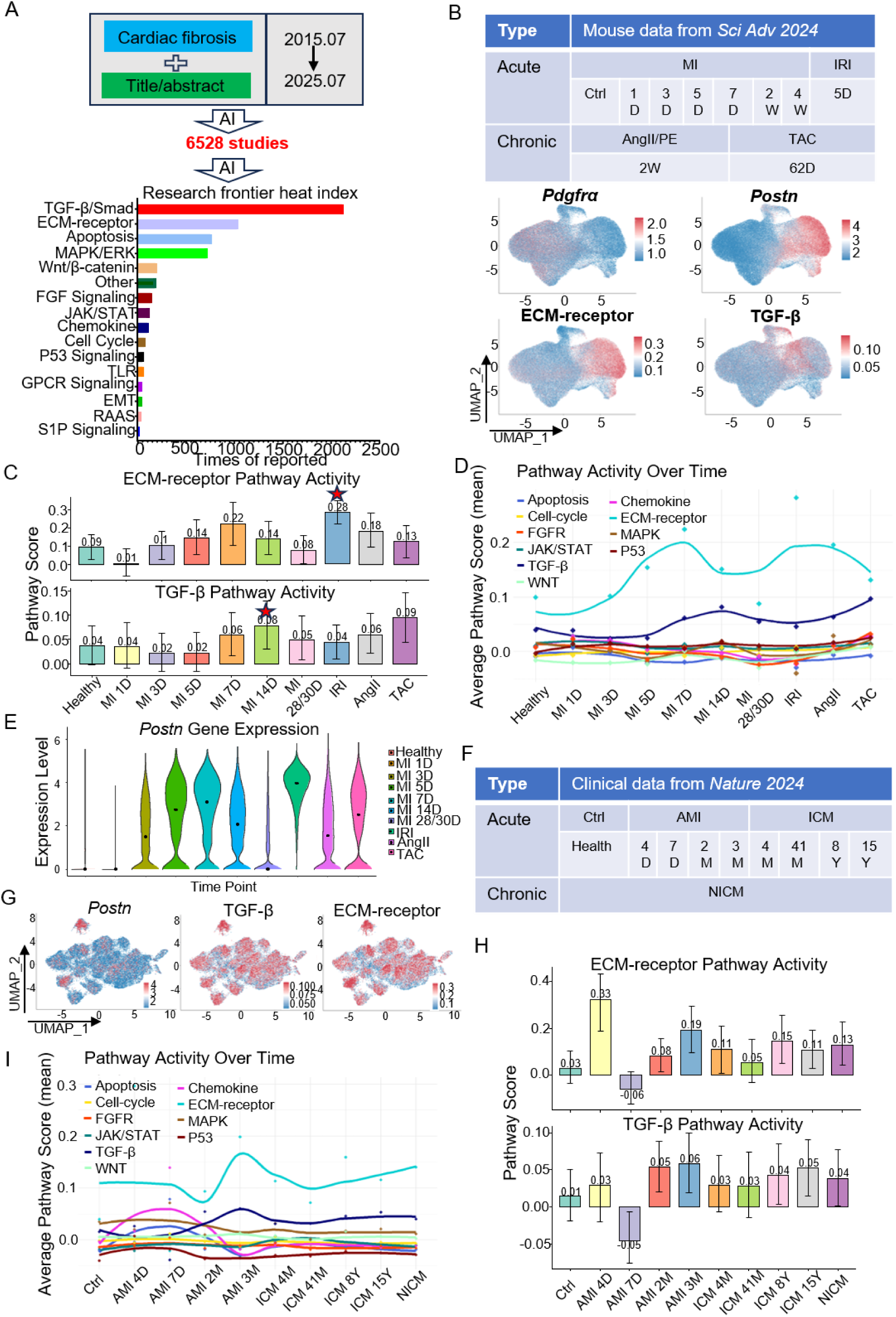
Temporal dynamics of core fibrosis-related signaling pathways across cardiac disease progression. A, AI-based literature mining of 6,528 publications on cardiac fibrosis (past decade) identified the core KEGG signaling pathways implicated in cardiac fibrosis. B, (above), Single-cell transcriptomic data of cardiac fibroblasts from diverse injury models (MI, IRI, AngII/PE, TAC) and injury stages (MI: 0, 1, 3, 5, 7, 14, 28 days) were analyzed for these pathways,(bottom): Spatial expression distribution of fibroblast/myofibroblast markers and ECM-receptor/TGF-β signaling pathways. C, Enrichment bar plots of the top two pathways (ECM and TGF-β) across injury models and stages. D, Dynamic enrichment curves of the top 10 pathways across injury models and stages. E, Expression kinetics of the myofibroblast marker *Postn* across injury models and stages. F, Pathway enrichment analysis of aggregated human clinical datasets. G, Spatial co-expression of myofibroblast markers with TGF-β and ECM-receptor pathways in clinical samples. H, Enrichment bar plots of ECM and TGF-β pathways across clinical subtypes and disease stages. I, Dynamic enrichment curves of the top 10 pathways in clinical cohorts stratified by subtype and stage.

Since identical pathways may exhibit distinct activation patterns across different fibrotic diseases and disease stages, we performed dynamic enrichment analysis using single-cell transcriptomic data from cardiac fibroblasts across multiple disease contexts^14,27,28^. We integrated data from murine acute injury models (MI; ischemia-reperfusion injury, IRI) and chronic fibrosis models (transverse aortic constriction, TAC; angiotensin II/phenylephrine, AngII/PE). Given the time-sensitive nature of MI as an acute injury model, the study included data from seven time points (0, 1, 3, 5, 7 days, 2 and 4 weeks post-MI; Figure 1B), making it an ideal resource for addressing our scientific questions in animal models. Spatial analysis revealed that *Pdgfrα* (a quiescent fibroblast marker^13^) and *Postn* (an activated myofibroblast marker^13^) exhibited mutual exclusivity (Figure 1B, lower panel), consistent with the established paradigm that cardiac fibrosis primarily involves fibroblast-to-myofibroblast transition^35^. Furthermore, ECM-receptor pathway activity (KEGG ID: hsa04512) strongly colocalized with *Postn*-high cells, while TGF-β pathway activity was predominantly confined to *Postn*^+^ and ECM-receptor^+^ cell populations (Figure 1B, lower panel). These findings suggest that fibroblast activation state, as indicated by *Postn* expression, specifically determines excessive ECM accumulation, whereas quiescent fibroblasts contribute minimally to pathological ECM deposition.

We then analyzed the enrichment of TGF-β and ECM pathways across different models and pathological stages (Figure 1C). The TGF-β pathway demonstrated higher enrichment in chronic models (TAC, AngII/PE) compared to acute models (MI, IRI). TAC day 62, representing advanced chronic fibrosis, exhibited the highest TGF-β enrichment, suggesting that TGF-β signaling constitutes a persistently activated pathway in chronic fibrosis and represents a conserved therapeutic target. In contrast, TGF-β pathway activity in MI showed notable fluctuations, peaking at two weeks post-injury before declining. Remarkably, ECM pathway activity peaked as early as one week post-MI, preceding the TGF-β maximum. This temporal dissociation indicates that ECM accumulation may occur independently of canonical TGF-β signaling.

Dynamic profiling of eight additional pathways revealed that TGF-β and ECM pathways were not only the most extensively studied but also exhibited the highest enrichment scores (Figure 1D).

Analysis of *Postn* expression, reflecting myofibroblast abundance and activation state, demonstrated peak expression around one week post-MI with subsequent decline to baseline levels by four weeks, consistent with previous report^14^ (Figure 1E). This temporal pattern suggests that *Postn*^+^ cells may not be optimal for investigating late-stage MI fibrosis mechanisms. Conversely, *Postn* expression remained elevated even at TAC day 62, indicating potential differences in terminal effector cell populations between acute and chronic fibrosis. These findings highlight significant mechanistic differences in myofibroblast activation between chronic and acute cardiac fibrosis, emphasizing the importance of precision medicine approaches in advanced fibrotic diseases.

To validate our findings, we analyzed pathway enrichment using clinical datasets from patients with acute and chronic cardiac fibrosis^14^, across various disease stages (Figure 1F). UMAP clustering analysis of human samples revealed that *POSTN* exhibited clear cell-type specificity, whereas ECM and TGF-β pathway activities showed broader distribution across fibroblast subpopulations (Figure 1G). Importantly, the highest activities of both pathways were observed in *POSTN^+^* cells, consistent with murine data. These findings suggest that in clinical contexts, ECM dysregulation may involve a broader fibroblast population beyond classical myofibroblasts, potentially providing new therapeutic insights for clinical antifibrotic strategies targeting the entire fibroblast population rather than the relatively scarce myofibroblast subset. This observation may also reflect species-specific differences in ECM-producing cell populations between mouse models and human pathology.

Temporal analysis of clinical samples confirmed that TGF-β and ECM pathways exhibited the highest enrichment scores (Figure 1H and 1I), consistent with murine data. ECM pathway activation preceded TGF-β activation and closely aligned with *Postn* expression kinetics (Figure S1B), reinforcing the tight coupling between early ECM initiation and myofibroblast activation. Among other pathways, the Chemokine pathway demonstrated higher activity than TGF-β during early-to-intermediate stages, suggesting its potential as a target for early-phase clinical intervention.

### Sequential activation of JAK-STAT and TGF-β pathways during cardiac fibrosis progression in MI

Our previous study demonstrated that genetic or pharmacological inhibition of JAK signaling attenuates angiotensin II/phenylephrine-induced cardiac fibrosis^29^. To enable precision therapeutic targeting, we conducted a comprehensive spatiotemporal analysis of JAK-STAT activation across different cardiac disease contexts. Analysis of clinical datasets and mouse models revealed distinct patterns of JAK-STAT pathway activation. In acute myocardial infarction, we observed robust early JAK-STAT pathway activation followed by a return to baseline levels (Figure 2A and 2B). In contrast, chronic fibrosis models exhibited sustained pathway activation, suggesting a critical role in progressive fibrotic remodeling. Initial analysis of murine MI models appeared to show incomplete recapitulation of the clinical pattern. However, further investigation revealed that only 88 of 254 core *Jak-Stat* genes were detected in the original dataset (detection rate: 34.6%), indicating potential technical limitations. Re-analysis focusing specifically on core JAK-STAT pathway components (e.g., *Jak1*, *Stat3*) demonstrated early upregulation consistent with clinical observations (Figure 2C).

**Figure 2.**
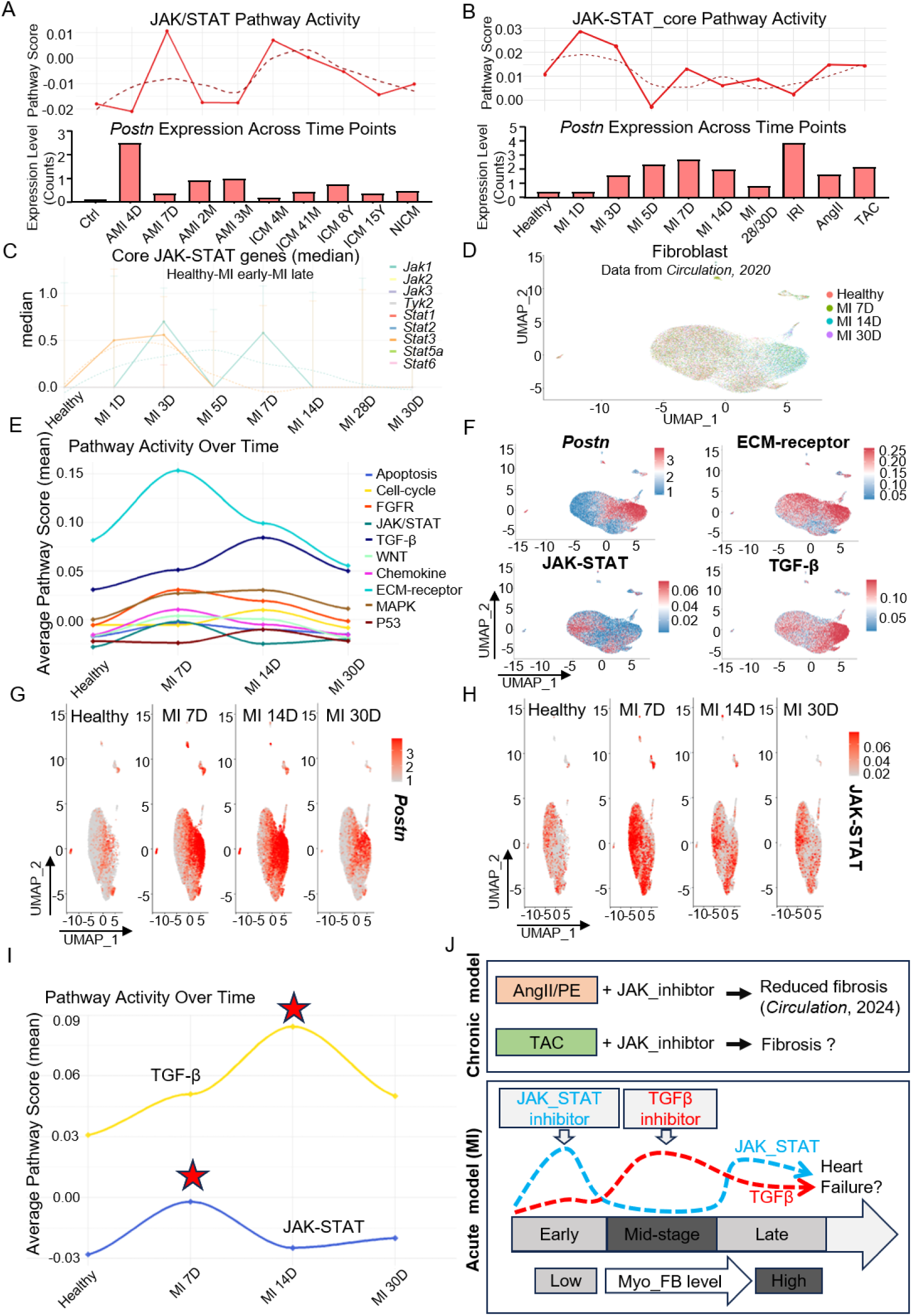
Sequential activation of JAK-STAT and TGF-β pathways during cardiac fibrosis progression. A, Temporal alignment of JAK-STAT pathway enrichment with fibroblast activation level timelines in clinical cohorts. B, JAK-STAT enrichment versus fibroblast activation level timelines in experimental injury models. C, Expression dynamics of JAK-STAT core genes during early/mid/late MI phases. D, UMAP plot of fibroblast across MI phases (0D, 7D, 14D and 30D). E, Dynamic pathway enrichment curves (top 10) across MI phases (0D, 7D, 14D and 30D). F, Spatial mapping of myofibroblast markers with ECM_receptor, TGF-β, and JAK-STAT pathways. G, Spatiotemporal expression of the activation marker *Postn* across four MI phases. H, JAK-STAT pathway activity across MI phases (spatiotemporal). I, JAK-STAT and TGF-β signaling pathways exhibit temporally staggered activation during cardiac fibrosis progression. J, Therapeutic model: “JAK-STAT inhibitor (early) + TGFB inhibitor (mid)” combinatorial regimen.

To minimize batch effects and normalization artifacts, we validated these findings using an independent dataset^27^ covering MI time points at days 0, 7, 14, and 30 (Figure 2D). This analysis confirmed dynamic changes in myofibroblast populations (Figure S2A) and temporal activation patterns across 10 pathways-including TGF-β, ECM-receptor, and apoptosis-consistent with our previous findings. Notably, the JAK-STAT pathway exhibited a distinct early peak (Figure 2E), coinciding with ECM-receptor activation and preceding TGF-β maximum, suggesting that JAK-STAT may influence ECM remodeling prior to TGF-β activation.

Spatial analysis demonstrated that both ECM and TGF-β signaling strongly colocalized with *Postn*^+^ cell subpopulations (Figure 2F), indicating robust association between fibroblast activation and these pathways in MI. Interestingly, JAK-STAT activation showed poor spatial correlation with *Postn*^+^ cells (Figure 2A and 2B), suggesting that JAK-STAT activation may not synchronize with *Postn* expression. To test this hypothesis, we evaluated UMAP projections of *Postn* expression and JAK-STAT pathway enrichment across four time points. *Postn* expression increased at week 1, peaked at week 2, then declined (Figure 2G), whereas JAK-STAT activation peaked at week 1 and subsequently decreased (Figure 2H and 2I; Figure S2B and S2C)-consistent with both clinical and murine datasets. These findings indicate that JAK-STAT activation precedes *Postn* expression in murine MI, supporting its potential role in promoting myofibroblast activation and fibrosis initiation.

Based on these comprehensive analyses, we propose the following therapeutic model (Figure 2J): First, given sustained JAK-STAT activation in chronic cardiac injury and its role in fibroblast activation, genetic ablation of JAK-STAT signaling in chronic models (e.g., AngII/PE) ameliorates fibrosis and cardiac hypertrophy^29^. Its therapeutic potential in other chronic models, such as TAC, warrants further investigation. Second, in acute MI, early JAK-STAT activation suggests that its inhibition may attenuate cardiac fibrosis development. Fibroblast-specific *Jak/Stat* knockout could provide therapeutic benefit, although the early activation peak may limit the efficacy of targeting JAK-STAT specifically in activated myofibroblasts. Third, leveraging the temporal dissociation between JAK-STAT (early phase) and TGF-β (intermediate phase) activation, we propose a “stage-specific dual-pathway inhibition” strategy: JAK-STAT inhibition during the early phase (days 1-8 post-MI) followed by TGF-β inhibition during the intermediate phase (days 6-14). This approach may overcome the temporal limitations of single-target inhibition (e.g., TGF-β peaks after scar formation is established) while minimizing adverse effects associated with prolonged JAK-STAT or TGF-β suppression.

### Myofibroblast-specific *Jak1/2* deletion attenuates cardiac fibrosis and hypertrophy in pressure overload-induced injury

Given the upregulation of JAK-STAT signaling observed in both AngII/PE and TAC models, we hypothesized that JAK-STAT inhibition could ameliorate chronic cardiac fibrosis. While our previous studies demonstrated that JAK-STAT pathway inhibition alleviates cardiac fibrosis and hypertrophy in the AngII/PE model^29^, we sought to determine whether JAK-STAT inhibition confers similar protective effects in the TAC model and to elucidate the underlying mechanisms.

We generated four myofibroblast-specific knockout mouse models using the Cre-loxP system (Figure 3A): control (*Postn^MCM/+^; Rosa26^NG/wt^*), *Jak1* knockout (*Postn^MCM/+^; Rosa26^NG/wt^; Jak^fl/fl^*), *Jak2* knockout (*Postn^MCM/+^; Rosa26^NG/wt^; Jak2^fl/fl^*), and *Jak1/Jak2* double knockout (*Postn^MCM/+^; Rosa26^NG/wt^; Jak1^fl/fl^; Jak2^fl/fl^*). Gene recombination was induced by intraperitoneal tamoxifen (TAM) injection, and cardiac function and structure were assessed two weeks post-TAC surgery (Figure 3B). Histological analysis revealed significantly attenuated cardiac fibrosis in all knockout groups compared to controls (Figures 3C and 3D), confirming the anti-fibrotic effects of *Jak1/2* deletion. Assessment of cardiac hypertrophy demonstrated significant reductions in the ventricle weight-to-body weight ratio (VW/BW) in the *Jak1* single-knockout group (Figure 3E), while WGA staining revealed decreased cardiomyocyte cross-sectional area in all knockout groups (Figures 3F and 3G), indicating amelioration of pressure overload-induced myocardial hypertrophy.

**Figure 3.**
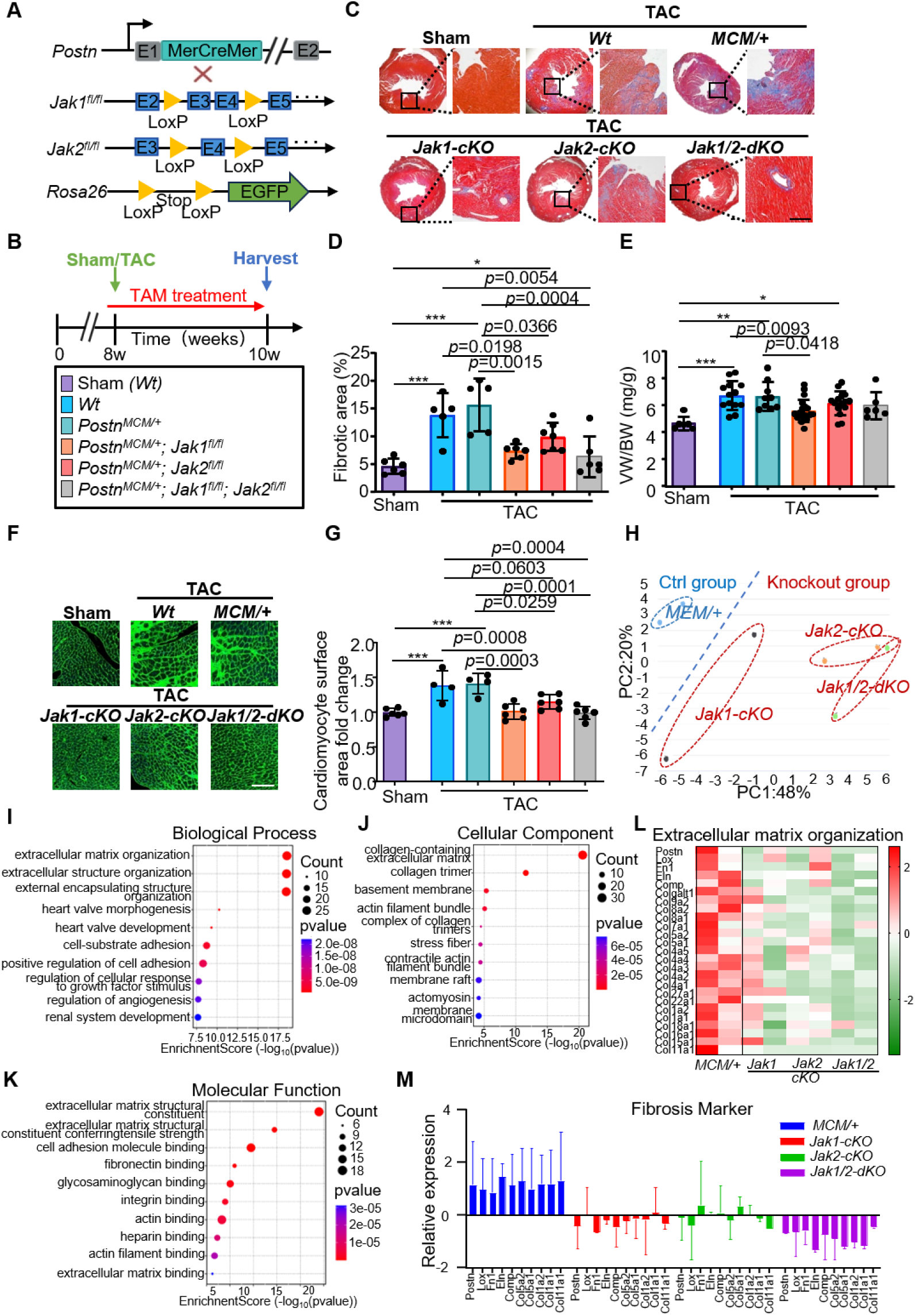
Myofibroblast-specific *Jak1/2* deletion attenuates cardiac fibrosis and hypertrophy in pressure overload-induced injury. A, Schematic representation of different mouse lines used, including the *Postn* genetic locus containing a tamoxifen-regulated MCM cDNA cassette inserted into exon 1 (E1), which was crossed with *Jak1-* and/or *Jak2-loxP*–containing gene-targeted lines, along with the *Rosa26* reporter allele(*R26^EGFP^*). B, Schematic representation of the experimental design. C, Representative Masson’s trichrome staining of cardiac tissue sections of Sham and TAC mice. Scale bar=250 μm. D, Quantification of fibrotic area (%) in each group. E, Ventricle weight–to–body weight ratio (VW/BW) following TAC administration. n=6: 13: 9: 21: 15: 6. F and G, Representative images of wheat germ agglutinin staining in cardiac tissue and quantification of cardiomyocyte surface area. Scale bar=150 μm. n=6: 4: 4: 6: 6: 6. H. PCA reveals distinct clustering of transcriptomic profiles in myofibroblasts under different gene knockout group. I through K. Gene Ontology (GO) enrichment analysis of downregulated genes in *Jak1/2*-knockout myofibroblasts: **(I)** biological processes, **(J)** cellular components, and **(K)** molecular functions. The most significantly enriched terms are highlighted based on –log_10_(*P* value). L. Heatmap of differentially expressed genes involved in extracellular matrix (ECM) organization, demonstrating altered transcriptional regulation in response to genetic perturbation. M. Relative expression levels of key fibrosis-related marker genes. \**P*<0.05; \*\**P*<0.01; \*\*\**P*<0.001 vs the sham group using 1-way ANOVA followed by Tukey multicomparisons test. Individual data are presented as aligned dot plots, with the mean and SD. *Postn*, *Periostin*; TAC, transverse aortic constriction.

Echocardiographic evaluation revealed that the 2-week TAC protocol did not induce detectable alterations in global systolic function, as assessed by ejection fraction (EF) and fractional shortening (FS; Figure S3A). Similarly, diastolic functional parameters, including isovolumic relaxation time (IVRT), mitral inflow E/A ratio, and E/e’ ratio, remained unchanged (Figure S3B). Morphologically, left ventricular chamber dimensions (LVID), posterior wall thickness (LVPW), and interventricular septal thickness (IVS) showed no significant differences (Figure S3A).

To validate the molecular basis of JAK1/2 inhibition’s ameliorative effects on cardiac fibrosis, we performed bulk RNA-seq on *Postn*⁺ myofibroblasts sorted from mice two weeks post-TAC (Figure S3C). Principal component analysis (PCA) demonstrated clear separation between the *Jak1-cKO*, *Jak2-cKO*, and *Jak1/Jak2-dKO* groups compared to controls (Figure 3H). The double-knockout group exhibited greater divergence from controls than single-knockout groups, suggesting more pronounced molecular alterations. Gene Ontology enrichment analysis of downregulated genes revealed significant associations with fibrosis-related pathways: Biological Process (BP) terms included ECM organization and extracellular structure organization (Figure 3I); Cellular Component (CC) terms encompassed collagen-containing ECM and collagen trimer (Figure 3J); and Molecular Function (MF) terms were enriched for ECM and collagen-related functions (Figure 3K). Normalized expression analysis of key ECM genes (Figure 3L) and core fibrotic markers (Figure 3M) consistently demonstrated that both single and double *Jak1/2* knockouts significantly suppressed ECM-related pathways and fibrotic markers compared to controls, with the double-knockout group showing the most pronounced inhibition. These findings demonstrate that fibroblast-specific *Jak1/2* knockout significantly attenuates cardiac fibrosis in chronic cardiac injury.

### Single-cell RNA sequencing identifies JAK1/2-responsive pro-fibrotic myofibroblast subpopulations in pressure overload

While bulk RNA sequencing demonstrated that *Jak1/2* knockout downregulates ECM and fibrosis-related gene expression, the underlying mechanisms governing myofibroblast differentiation and functional heterogeneity require high-resolution validation. To address this, we performed single-cell RNA sequencing (scRNA-seq) on cardiac myofibroblasts (*Postn*⁺) from mice subjected to two weeks of TAC, providing the first systematic characterization of myofibroblast heterogeneity under pathological conditions and its dependence on JAK-STAT signaling.

To investigate myofibroblast clustering patterns and identify subpopulations with enhanced ECM secretion within the *Postn⁺* population, we first analyzed *Postn*⁺ cells from control mice. UMAP dimensionality reduction identified seven distinct subpopulations (Figure 4A). Notably, ECM secretion capacity varied among myofibroblast subpopulations, with Cluster 1 exhibiting markedly elevated expression of ECM-related genes and significantly higher ECM scores compared to other clusters (Figure 4B through 4D). Interestingly, while UMAP analysis suggested an overall correlation between *Postn* expression and ECM secretion, *Postn* expression in Cluster 1 was not the highest (Figure S4A), indicating dissociation between *Postn* expression and ECM secretion capacity in myofibroblasts.

**Figure 4.**
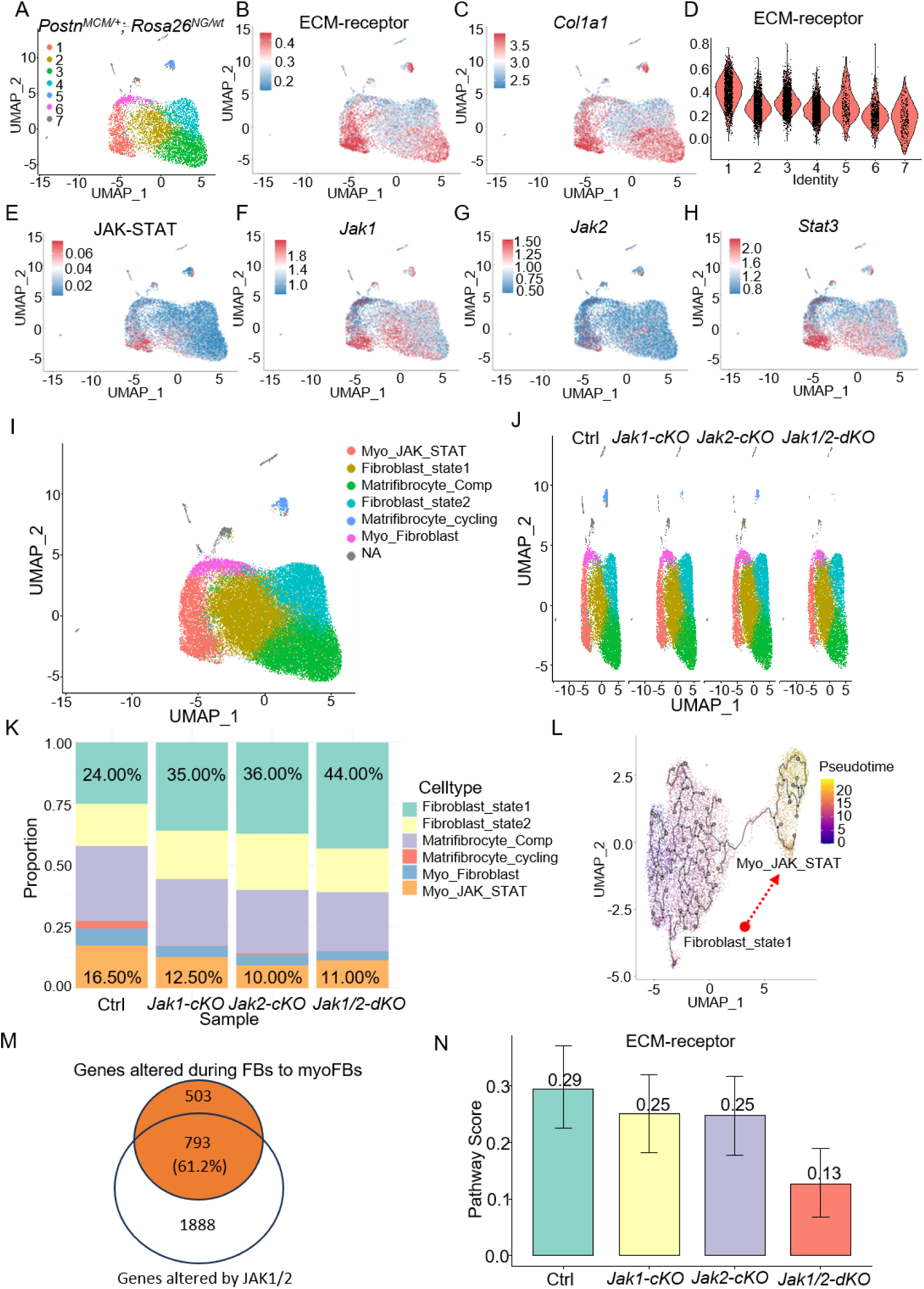
Single-cell RNA sequencing identifies JAK1/2-responsive pro-fibrotic myofibroblast subpopulations in pressure overload. A, UMAP plot of myofibroblasts colored by sample identity, ctrl group (*Postn^MCM/+^; Rosa26^NG/wt^*). B and C, UMAP plot of ECM-receptor and *Col1a1*, ctrl group (*Postn^MCM/+^; Rosa26^NG/wt^*). D, ECM-receptor expression in myofibroblasts by clusters as violin plot. E through H, UMAP plot of *Jak/Stat* family gene expression in myofibroblasts, ctrl group (*Postn^MCM/+^; Rosa26^NG/wt^*). I, UMAP plot of myofibroblasts colored by clusters identity. J. UMAP plot showing cardiac fibroblast heterogeneity across groups. K. Percentage of cell population in the control and knockout groups. L, Pseudotemporal trajectory analysis showed Fibroblast_state1 to Myo_JAK_STAT transition. M, Genes altered during Fibroblast_state1 to Myo_JAK_STAT (above circle), genes altered by *Jak1/2* (bottom circle). N, Pathway Score of ECM-receptor among four groups.

To determine whether JAK-STAT signaling correlates with the high ECM secretion observed in Cluster 1, we assessed JAK-STAT pathway activity. Remarkably, Cluster 1 exhibited both the highest JAK-STAT activation and the strongest ECM secretion (Figure 4E). Key pathway genes (*Jak1*, *Jak2*, *Stat3*) were significantly upregulated in this cluster (Figure 4F through 4H). Based on these distinctive molecular characteristics, we designated Cluster 1 as the “Myo_JAK_STAT” subpopulation-a pathogenic myofibroblast subset characterized by high ECM secretion and enhanced JAK-STAT activation.

To elucidate the impact of *Jak1/2* deletion on myofibroblast heterogeneity, we compared scRNA-seq data from Ctrl, *Jak1*-cKO, *Jak2*-cKO, and *Jak1/2*-dKO groups (approximately 40,000 cells total). Cell annotation identified seven subpopulations (Figure 4I and 4J): Matrifibrocyte_Comp (high Comp expression, representing the largest proportion, Figure S4B), Fibroblast_state1 (high *Pdgfrα*/*Tcf21*, quiescent fibroblasts, Figure S4C), Fibroblast_state2 (high *Tmem100*/*Cd248*, quiescent characteristics), Myo_JAK_STAT (high mitochondrial metabolic gene expression, enhanced ECM secretion, elevated mitochondrial metabolic activity, and JAK-STAT enrichment, Figure S4D), Matrifibrocyte_cycling (high proliferation genes, e.g., *Mki67*/*Top2a*, Figure S4E), and Myo_Fibroblast (high *Cmss1*/*Lars2*).

Population analysis revealed that Matrifibrocyte_cycling cells were markedly reduced in *Jak* knockout groups and nearly absent in the double-knockout group (Figure 4K), indicating that *Jak*/*Stat* knockout suppresses proliferation in the most proliferative myofibroblast subpopulation. Additionally, *Jak1/2* knockout groups exhibited a significant increase in Fibroblast_state1 (the subpopulation with lowest ECM secretion) and a substantial decrease in Myo_JAK_STAT (the subpopulation with highest ECM secretion; Figure 4K). Under injury conditions, the former likely differentiates into the latter. This provides direct evidence that *Jak* knockout impedes myofibroblast differentiation into the most fibrogenic subpopulation (Myo_JAK_STAT), leading to downregulation of core pro-fibrotic genes (Figure S4F) and representing a key mechanism by which *Jak* knockout ameliorates cardiac fibrosis.

Pseudotime analysis simulating the differentiation process in controls (Figure 4L) identified 1,296 genes (*p* < 0.001) significantly regulating fibroblast differentiation into Myo_JAK_STAT. These genes were enriched in pathways related to ECM assembly and mitochondrial metabolism (Figure S4G). Following *Jak1/2* knockout, 61.2% (793/1,296) of these key differentiation genes were significantly perturbed (Figure 4M), confirming that JAK signaling serves as a central driver of this differentiation process. Consequently, ECM-receptor pathway activity was downregulated in a dose-dependent manner in *Jak*-cKO groups (moderate reduction in single knockouts, substantial reduction in double knockout; Figure 4N), consistent with bulk RNA-seq findings (Figure 3H through 3M).

### Myofibroblast-specific *Jak1/2* deletion does not ameliorate cardiac dysfunction and fibrosis following myocardial infarction

In contrast to the chronic cardiac injury induced by TAC, acute ischemic injury from MI triggers intense inflammatory responses and rapid tissue remodeling^8,36,37^. Our analysis demonstrated that myofibroblasts proliferate rapidly around day 7 to form scar tissue and begin to regress after two weeks post-MI. Therefore, precise timing is critical for JAK-based therapeutic interventions in the MI model. We hypothesized that although *Postn*-mediated myofibroblast *Jak* knockout proved effective in the TAC model, it might be ineffective in MI (Figure 5A), partly because peak JAK-STAT expression occurs earlier than peak *Postn* expression (Figure 2G and 2H).

**Figure 5.**
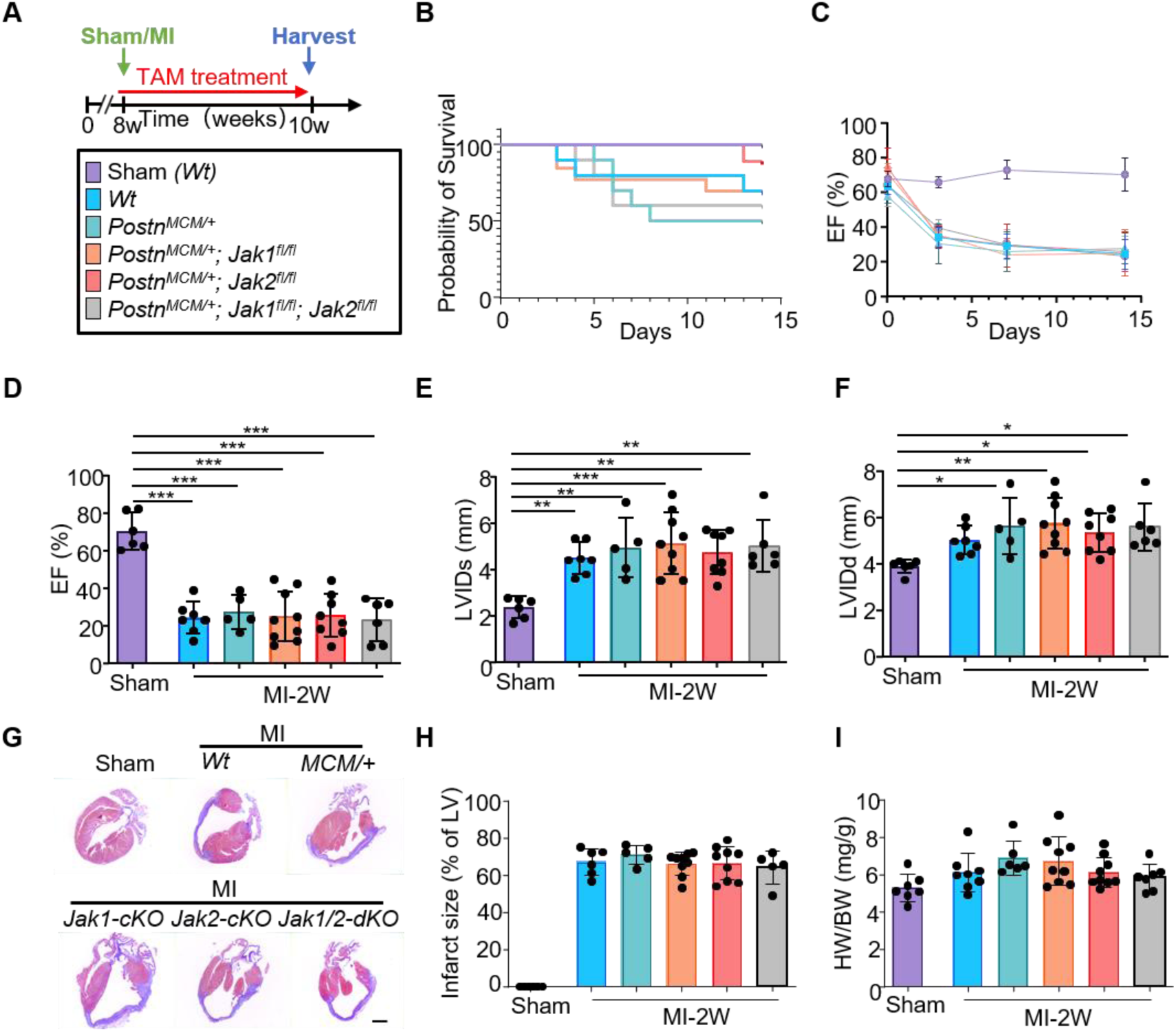
Myofibroblast-specific *Jak1/2* deletion does not ameliorate cardiac dysfunction and fibrosis following myocardial infarction. A, Schematic representation of the experimental design. Mice were subjected to MI surgery at 8 weeks of age, harvested at 10 weeks, and given TAM treatment by intraperitoneal injection within two weeks to induce gene knockout. B, Survival plot in days following MI injury for the five genotypes of mice shown (n=10 for *Wt*, *Postn^MCM/+^* and *Postn^MCM/+^; Jak1^fl/fl^; Jak2^fl/fl^* separately, n=13 for *Postn^MCM/+^; Jak1^fl/fl^* and n=9 for *Postn^MCM/+^; Jak2^fl/fl^*). C, Representation showing the dynamics of the LV ejection fraction (EF) along infarction. Cardiac function was evaluated at day 0, 3, 7 and 14 after MI. D through F, Quantitative evaluation of EF (%), LV systolic and diastolic internal diameter (LVIDs and LVIDd) in sham mice compared with MI mice. n=6: 7: 5: 9: 8: 6. G, Representative Masson’s trichrome staining of longitudinal heart sections from Sham and MI mice: Wild-type (Wt), *Postn^MCM/+^* (control for Cre expression), *Jak1-cKO* (*Postn^MCM/+^; Jak1^fl/fl^*), *Jak2-cKO* (*Postn^MCM/+^; Jak2^fl/fl^*), and *Jak1/2-dKO* (*Postn^MCM/+^; Jak1^fl/fl^; Jak2^fl/fl^*). Scale bar=1 mm. H, Quantification of the infarct size (% of LV). n=6: 6: 5: 9: 9: 5. I, Heart weight–to–body weight ratio (HW/BW) following MI administration. n=7: 8: 6: 9: 8: 7. \**P*<0.05; \*\**P*<0.01; \*\*\**P*<0.001 vs the sham group using 1-way ANOVA followed by Tukey multicomparisons test. Individual data are presented as aligned dot plots, with the mean and SD. MI, myocardial infarction; TAM, tamoxifen; *Postn*, *Periostin*.

Kaplan-Meier survival analysis revealed that myofibroblast-specific knockout of *Jak1/2* did not improve post-MI survival compared to controls (Figure 5B). Serial echocardiographic assessment indicated no significant improvement in left ventricular systolic function parameters (EF and FS) in any knockout group (Figure 5C through 5F; Figure S5), suggesting that inhibiting JAK1/2 signaling in myofibroblasts does not mitigate post-MI cardiac dysfunction. Masson’s trichrome staining revealed no significant differences in infarct size between knockout and control groups (Figure 5G and 5H), indicating that *Jak1/2* deletion does not affect the extent of infarction-induced fibrotic remodeling. The heart weight-to-body weight ratio (HW/BW), an indicator of hypertrophic response, also showed no significant differences among groups (Figure 5I).

These data indicate that since the peak of *Postn*^+^ myofibroblasts occurs after the most active phase of the JAK-STAT pathway, *Postn^MCM/+^*-mediated myofibroblast-specific *Jak1/2* knockout has limited efficacy on post-MI fibrosis and cardiac dysfunction.

### Preventive genetic ablation of *Jak1/2* in cardiac fibroblasts attenuates post-MI fibrosis and adverse remodeling

We then test if inhibiting *Jaks* before their peak activity might ameliorate MI-induced cardiac fibrosis. *Pdgfrα*-creERT mice were crossed with *Jak1^fl/fl^* and *Jak2^fl/fl^* mice to generate fibroblast-specific *Jak1/2* knockout models (Figure 6A). The experimental timeline included gene knockout induction at 6-8 weeks of age, followed by left anterior descending coronary artery ligation at 9 weeks, with cardiac function and tissue remodeling assessed at two weeks post-MI (Figure 6B).

**Figure 6.**
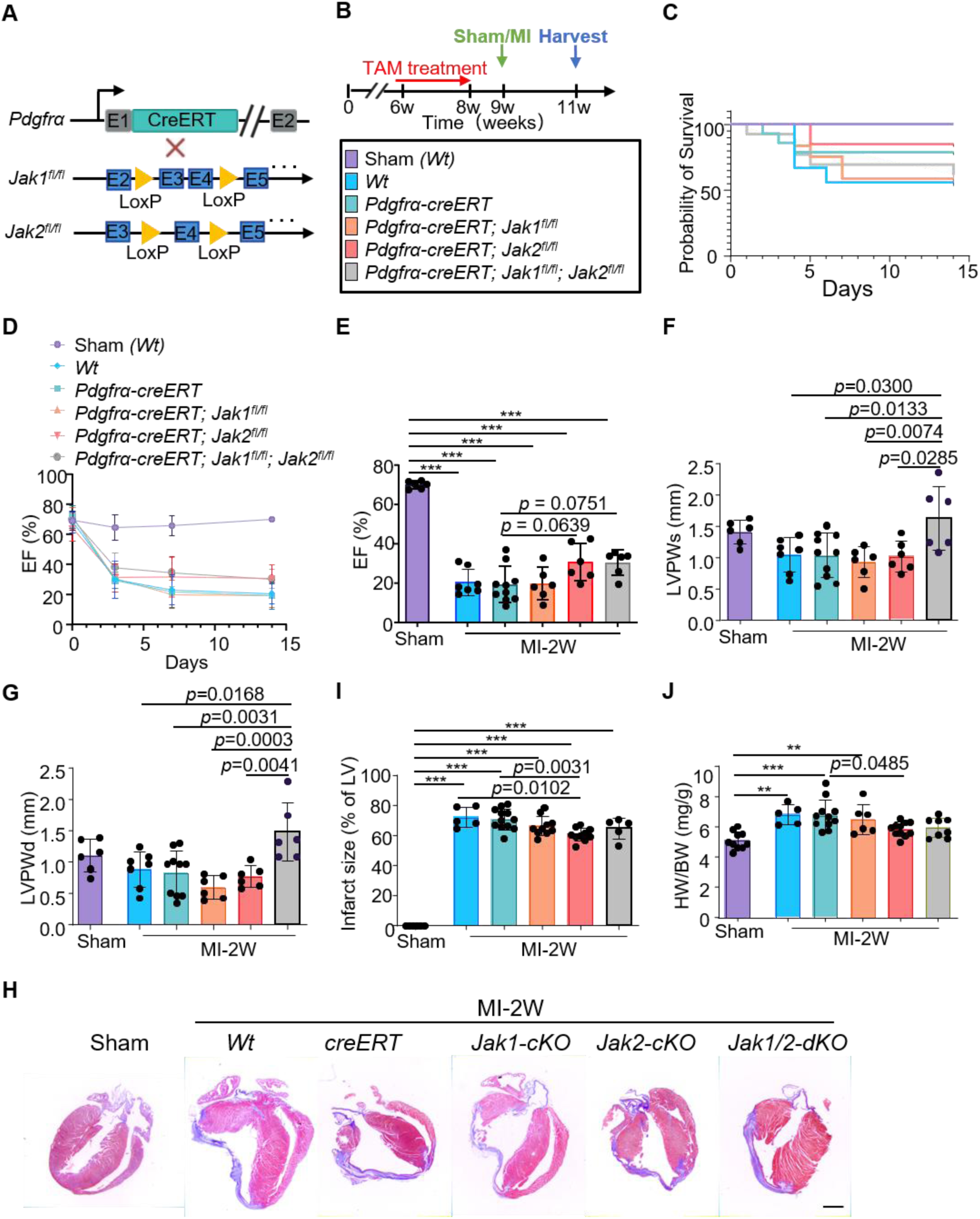
Preventive genetic ablation of *Jak1/2* in cardiac fibroblasts attenuates post-MI fibrosis and adverse remodeling. A, Schematic representation of different mouse lines used, including the *Pdgfrα* genetic locus containing a TAM-regulated CreERT cDNA cassette inserted into exon 1 (E1), which was crossed with *Jak1-* and/or *Jak2-loxP*–containing gene-targeted lines. B, Schematic representation of the experimental design. Mice were administered TAM treatment via intraperitoneal injection from 6 to 8 weeks of age. MI surgery was conducted at 9 weeks, and mice were harvested at 11 weeks. C, Probability of survival of mice after MI. n=6: 9: 14: 12: 13: 13. D through G, Quantification of LVEF, LVPWs and LVPWd. n=6: 7: 10: 6: 6: 6. H, Representative Masson’s trichrome staining of longitudinal heart sections from Sham and MI mice: Wild-type (Wt), *Pdgfra-creERT* (control for Cre expression), *Jak1-cKO* (*Pdgfra-creERT; Jak1^fl/fl^*), *Jak2-cKO* (*Pdgfra-creERT; Jak2^fl/fl^*), and *Jak1/2-dKO* (*Pdgfra-creERT; Jak1^fl/fl^; Jak2^fl/fl^*). Scale bar=1 mm. I, Quantification of the infarct size (% of LV). n=6: 5: 12: 10: 10: 5. J, Heart weight–to–body weight ratio (HW/BW) following MI administration. n=10: 5: 11: 6: 11: 8. \*\**P*<0.01; \*\*\**P*<0.001 vs the sham group using 1-way ANOVA followed by Tukey multicomparisons test. Individual data are presented as aligned dot plots, with the mean and SD. MI, myocardial infarction; TAM, tamoxifen; LVEF, left ventricular ejection fraction; LVPWs and LVPWd, left ventricular posterior wall in systole and diastole.

Kaplan-Meier survival analysis revealed no significant differences between knockout and control groups (Figure 6C). Serial echocardiographic assessments at baseline and 3, 7, and 14 days post-MI demonstrated comparable EF values across all groups (Figure 6D and 6E; Figure S6). Notably, the double-knockout group exhibited attenuated thinning of the left ventricular posterior wall, indicating reduced adverse remodeling (Figure 6F and 6G). Most importantly, Masson’s trichrome staining revealed a significant reduction in infarct size in fibroblast-specific *Jak2* knockout mice compared to controls (Figure 6H and 6I). Furthermore, the heart weight-to-body weight (HW/BW) ratio was significantly decreased in the *Jak2* knockout group relative to control animals (Figure 6J).

Collectively, these findings demonstrate that while fibroblast-specific *Jak1/2* knockout does not ameliorate post-MI systolic dysfunction, it partially attenuates adverse cardiac remodeling and reduces fibrotic tissue deposition following myocardial infarction.

### Ruxolitinib treatment attenuates post-MI cardiac fibrosis and improves cardiac function

To evaluate the therapeutic potential of JAK1/2 inhibition in myocardial infarction, we administered the selective JAK1/2 inhibitor RUX prior to the observed JAK activity peak. To ensure adequate drug exposure, oral gavage was initiated one day before MI surgery and continued for 14 days, followed by cardiac tissue collection (Figure 7A). Kaplan-Meier survival analysis demonstrated no significant differences between treatment groups (Figure 7B).

**Figure 7.**
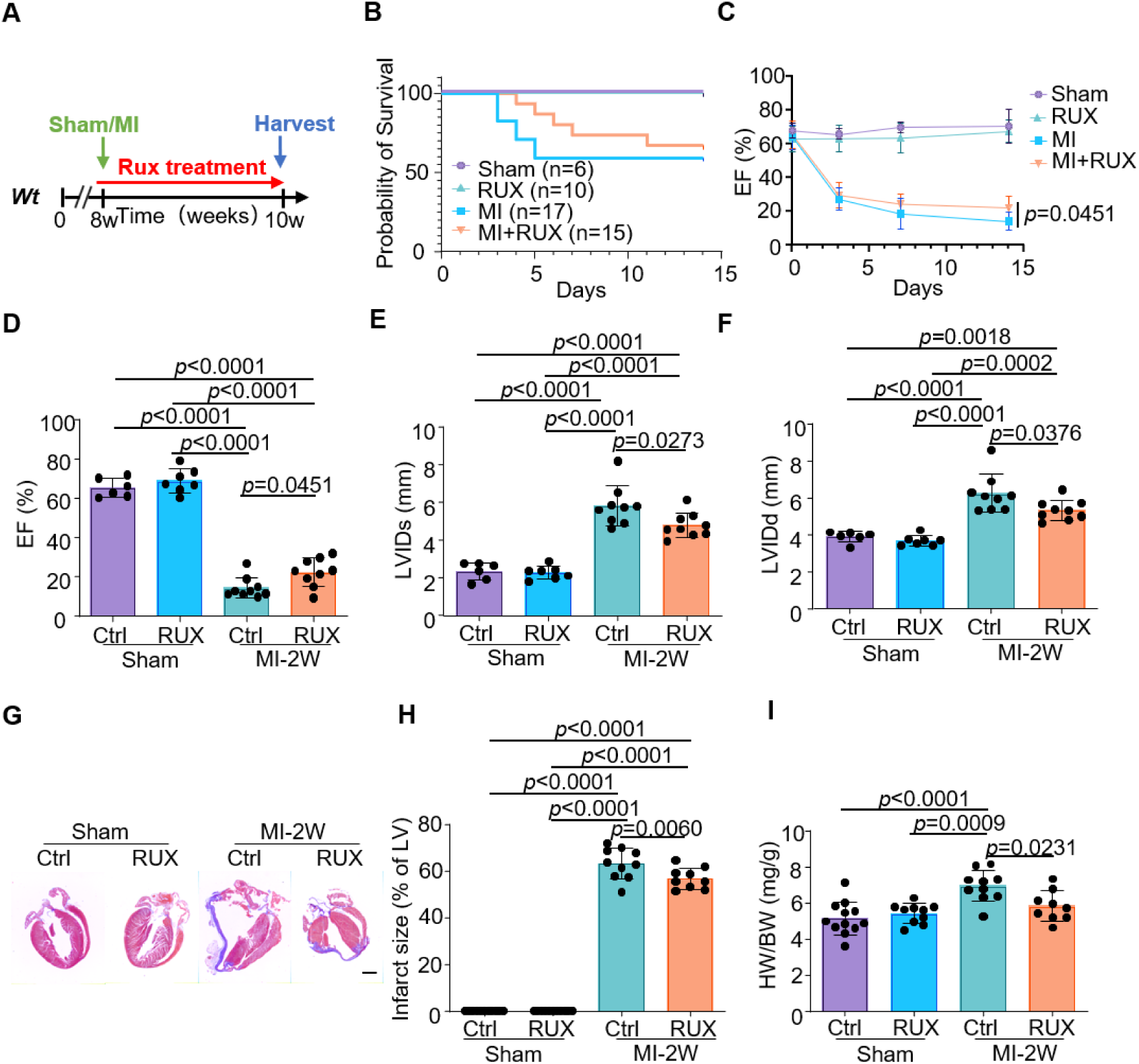
Ruxolitinib treatment attenuates post-MI cardiac fibrosis and improves cardiac function. A, Schematic representation of the experimental design. *Wt* mice were subjected to MI at 8 weeks, and treatment with Rux was administered via gavage concurrently with MI. The mice were harvested at 10 weeks. B, Kaplan-Meier survival curves in Sham, MI, Rux and MI+Rux mice. n=6: 10: 17: 15. C, Echocardiography to determine changes in LVEF at 0, 3, 7, and 14 days after MI. D through F, Echocardiography to measure EF (%), LVIDs and LVIDd 14 days post-MI, n=6: 7: 9: 9. G, Representative images of Masson’s trichrome staining of longitudinal heart sections from Sham, Rux, MI and MI+Rux mice. Scale bar=1 mm. H, Quantification of the infarct size (% of LV). n=11: 11: 10: 9. I, Heart weight–to–body weight ratio (HW/BW) following MI administration. n=12: 10: 10: 9. 1-way ANOVA was used followed by the Tukey multiple comparisons test. Individual data are presented as aligned dot plots, with the mean and SD. *Wt*, wild type; MI, myocardial infarction; Rux, ruxolitinib; EF, ejection fraction; LVIDs, left ventricular end-systolic internal diameter; LVIDd, left ventricular end-diastolic internal diameter.

Echocardiographic assessment revealed significantly improved EF in the RUX-treated group compared to the MI control group at day 14 post-infarction (Figure 7C and 7D; Figure S7). Additionally, RUX treatment resulted in improved left ventricular internal dimensions in both systole (LVIDs) and diastole (LVIDd) (Figure 7E and 7F). Masson’s trichrome staining demonstrated significantly reduced cardiac fibrosis in the RUX-treated group (Figure 7G and 7H), accompanied by a significantly lower heart weight-to-body weight (HW/BW) ratio (Figure 7I).

These results confirm that pre-emptive JAK inhibition represents an effective therapeutic strategy for cardiac fibrosis, highlighting the critical importance of therapeutic timing. Based on these observations, we developed a multi-pathway combination therapy guided by dynamic pathway enrichment profiles during MI progression. We selected the TGF-β signaling pathway for combination with JAK-STAT inhibition, given its prominent activity profile and availability of well-established inhibitors.

### Combined ruxolitinib and pirfenidone therapy attenuates MI-induced cardiac fibrosis and improves cardiac function

Based on the temporal activity profiles of both signaling pathways during MI progression, we designed a precisely timed combination treatment regimen (Figure 8A). Oral RUX was administered from one day before MI induction through day 8 post-MI to target peak JAK-STAT activity (day 7), while PFD, a TGF-β inhibitor, was administered from day 6 through day 14 post-MI to coincide with peak TGF-β pathway activation (day 14).

**Figure 8.**
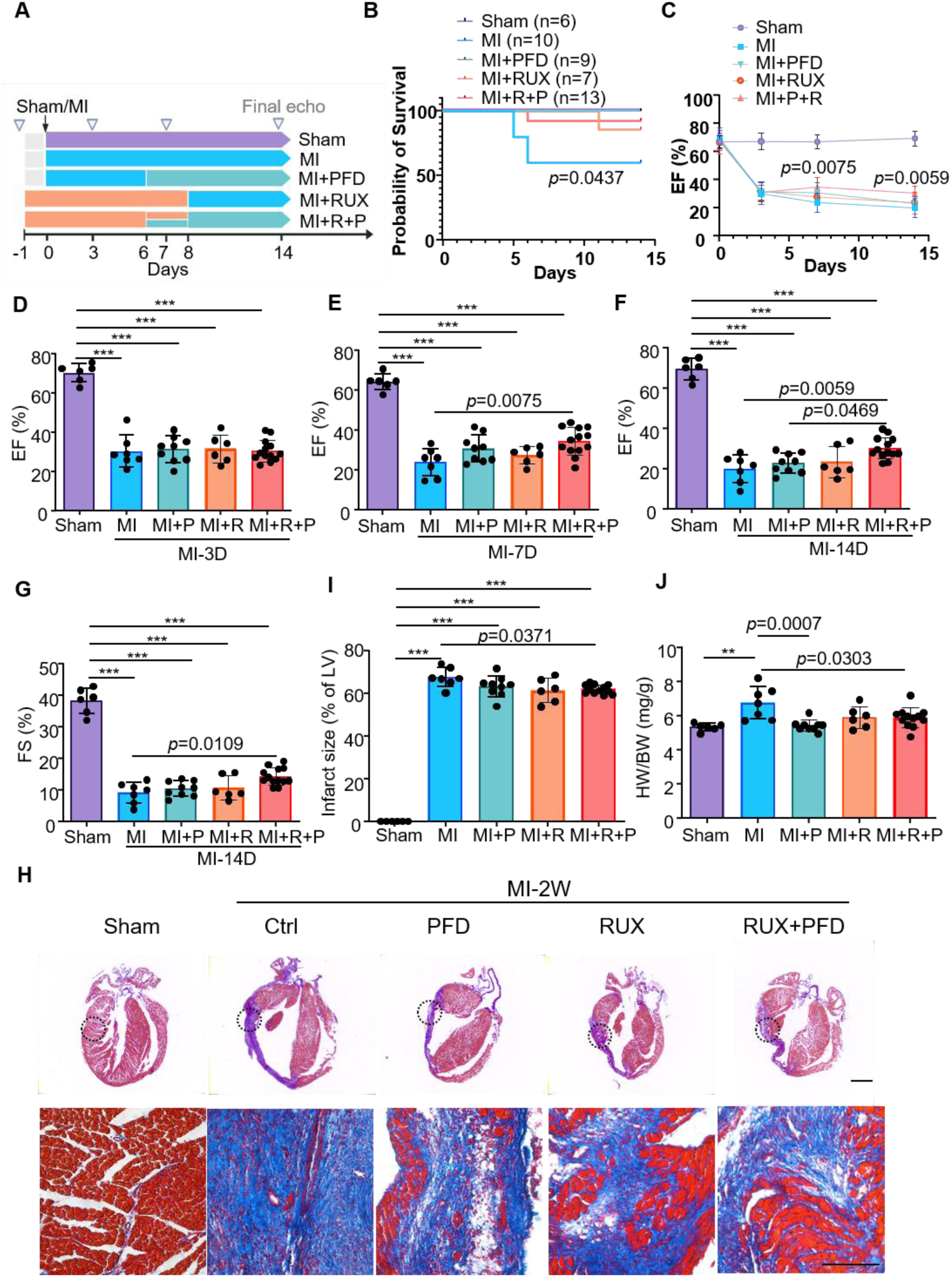
Combined ruxolitinib and pirfenidone therapy attenuates MI-induced cardiac fibrosis and improves cardiac function. A, Experimental design of pharmacological interventions in MI. Sham (n=6). MI control (MI with vehicle administration, n=10). MI+PFD (PFD monotherapy initiated on postoperative day 6 through endpoint, n=9). MI+RUX (RUX administered 1 day pre-MI through postoperative day 8, n=7). MI+RUX+PFD (combined RUX (−1 to 8d) and PFD (6 to 14d), n=13). B, Survival curves for the five treatment groups. C through G, Time-course of left ventricular ejection fraction, LVEF (%), and echocardiography measurements of LVEF (%), LVFS (%) at different time points. H, Representative images of Masson’s trichrome-stained tissue sections. Scale bars represent 1 mm (above) and 150 μm (bottom). I, Quantification of the infarct size (% of LV). n=6: 7: 9: 6: 12. J, Heart weight–to–body weight ratio (HW/BW) following MI administration. \*\**P*<0.01; \*\*\**P*<0.001 vs the sham group using 1-way ANOVA followed by Tukey multicomparisons test. Individual data are presented as aligned dot plots, with the mean and SD. MI, myocardial infarction; Rux, ruxolitinib; PFD, pirfenidone; EF, ejection fraction; FS, fraction shortening, LVIDs, left ventricular end-systolic internal diameter; LVIDd, left ventricular end-diastolic internal diameter.

Kaplan-Meier survival analysis demonstrated no significant differences among treatment groups (Figure 8B). We subsequently evaluated EF, a key indicator of heart failure severity (Figure 8C). At day 3 post-MI, no significant differences were observed between combination and monotherapy groups (Figure 8D). However, by day 7, the combination therapy group demonstrated significantly improved EF, indicating synergistic effects of concurrent JAK-STAT and TGF-β pathway inhibition (Figure 8E). At two weeks post-MI, the combination group maintained significantly superior EF and FS (Figure 8F and 8G; Figure S8), suggesting substantial preservation of cardiac function.

Masson’s trichrome staining and quantitative analysis revealed significantly reduced infarct size in the combination therapy group (Figure 8H and 8I). Morphologically, combination therapy significantly attenuated cardiac hypertrophy (Figure 8J). Notably, isolated islands of viable cardiomyocytes were observed within the infarct region, most prominently near the ligation site, further confirming that this therapeutic strategy improves both cardiac function and structural preservation following MI (Figure 8H).

## Discussion

Cardiovascular diseases account for one-third of global deaths and progress relentlessly through maladaptive remodeling driven by uncontrolled fibrosis^35,38–41^. This study provides the first comprehensive temporal mapping of fibrotic pathway activation following myocardial infarction and demonstrates that stage-specific dual-pathway inhibition significantly outperforms single-target interventions. Our findings suggest a shift from the current paradigm of anti-fibrotic therapy by demonstrating that intervention timing, rather than pathway selection alone, may be critical for determining therapeutic efficacy.

Recent advances in cardiac fibroblast research have enabled increasingly refined classifications of fibroblast subtypes through single-cell RNA sequencing^42–48^. While previous studies identified functionally distinct fibroblast subpopulations, high-resolution characterization of myofibroblast heterogeneity has remained elusive. Our single-cell analysis identified a highly pathogenic myofibroblast subpopulation (Myo_JAK_STAT) characterized by elevated ECM secretion and metabolic activity. The dramatic reduction of this subpopulation following JAK inhibition demonstrates that therapeutic efficacy depends on targeting specific cellular subsets rather than broad pathway suppression.

Importantly, comparative knockout studies revealed non-redundant roles of JAK1 and JAK2, indicating that complete JAK-STAT blockade is necessary to effectively disrupt fibroblast-to-myofibroblast differentiation. This cellular specificity explains why *Jak1/2* double knockout produced superior outcomes compared to individual knockouts, providing mechanistic insight into the superiority of dual JAK targeting.

The contrasting outcomes between *Pdgfrα-creERT* and *Postn^MCM/+^*-driven *Jak1/2* deletion underscore a critical principle: therapeutic windows are dictated by pathway activation kinetics, not cell-type specificity alone. Our pathway mapping revealed that JAK-STAT activity peaks before *Postn*^+^ cell expansion, explaining why *Postn^MCM/+^*-driven deletion occurs after the optimal intervention window. This temporal mismatch has important implications for clinical trial design, where intervention timing relative to disease onset may be more critical than previously recognized.

Current anti-fibrotic approaches, including angiotensin-converting enzyme (ACE) inhibitors, aldosterone antagonists, and statins, achieve only partial reverse remodeling through indirect suppression of fibroblast activation^49–53^. Substantial therapeutic gaps persist due to the lack of direct anti-fibrotic agents and dose-limiting side effects of existing therapies. The clinical availability of JAK inhibitors and TGF-β pathway modulators provides an immediate translational opportunity.

However, our findings suggest that current clinical trial approaches, which typically test single agents at fixed time points, may be fundamentally flawed^19^. Our kinetic analysis revealed that TGF-β activity peaks after initial scar formation, suggesting that TGF-β inhibition alone misses the early inflammatory-fibrotic transition. The synergistic effects of combining early JAK-STAT inhibition with mid-phase TGF-β blockade demonstrate that sequential polytherapy can overcome monotherapy limitations. We propose that mechanism-informed sequential regimens, guided by pathway activation biomarkers, represent a more rational therapeutic approach.

Our comparative analysis of signaling pathway signatures in human and murine fibroblasts reveals critical translational implications. The broader distribution of ECM-producing activity across human fibroblast populations, rather than the discrete *Postn*^+^ segregation observed in mice, suggests that clinical anti-fibrotic strategies may require targeting a wider cellular spectrum. This finding has implications for biomarker development and patient stratification in future clinical trials.

## Acknowledgements

This work was supported by the National Key R&D Program of China (2022YFA0806200), the Guangdong Natural Science Foundation (2024A1515013130), and the Science and Technology Program of Guangzhou (2024D03J0014). Dr. Molkentin and Dr. Huo were supported by grants from the National Institutes of Health (R01HL160765) and the American Heart Association (25CDA1447309), respectively.

## Nonstandard Abbreviations and Acronyms

ECM: extracellular matrix
HW/BW: heart weight-to-body weight
JAK1: Janus kinase 1
JAK2: Janus kinase 2
MI: myocardial infarction
PFD: pirfenidone
POSTN: periostin
RUX: ruxolitinib
TAC: transverse aortic constriction
TGF-β: transforming growth factor-β
VW/BW: ventricular weight-to-body weight

## Notes

### Competing Interest Statement

The authors have declared no competing interest.

### Summary of Updates

First, typographical errors in co-authors' names (Dr. Qiongjie Mi and Ningxin Gao) have been corrected. Second, the supplemental file containing one supplemental table and eight supplemental figures has been uploaded.

