## supplemental table 1 and supplemental figure 1-8 for "AI-based Predictive Signaling Pathway Profiling in Cardiac Fibrosis Suggests a Novel Combinatorial Treatment Strategy"

**SUPPLEMENTAL MATERIAL**

#### SUPPLEMENTAL TABLES:

**Table S1. Rank of Enrichment Frequencies in Cardiac Fibrosis-Related Signaling Pathways.**

| Pathway Group | Report Count |
|---|---|
| TGF-β/Smad | 2132 |
| ECM Receptor | 1041 |
| Apoptosis | 773 |
| MAPK/ERK | 726 |
| Wnt/β-catenin | 204 |
| Other | 189 |
| FGF Signaling | 155 |
| JAK-STAT | 124 |
| Chemokine | 113 |
| Cell Cycle | 80 |
| P53 Signaling | 70 |
| TLR | 67 |
| GPCR Signaling | 48 |
| EMT | 45 |
| RAAS | 38 |
| S1P Signaling | 22 |

### SUPPLEMENTAL FIGURES AND FIGURE LEGENDS:

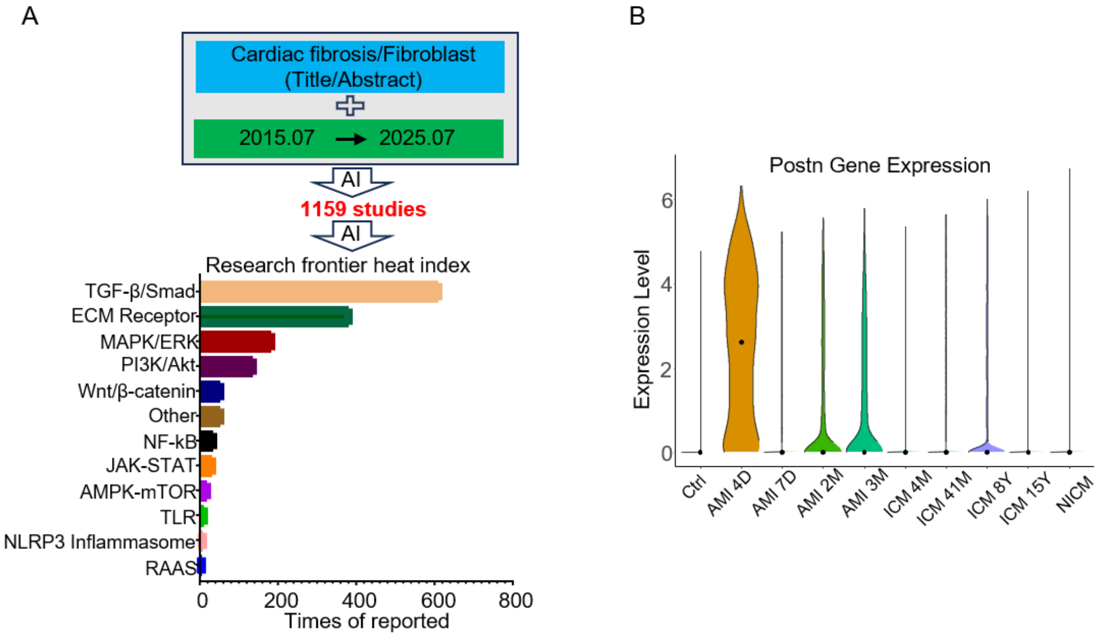


**Figure S1. Score Chart for Core Signaling Pathways in Fibroblasts.**

A, AI-based literature mining of 1,159 publications on fibroblast and cardiac fibrosis (past decade) identified the core KEGG signaling pathways implicated in cardiac fibrosis.

B, Expression of the myofibroblast marker *Postn* across clinical subtypes and disease stages.

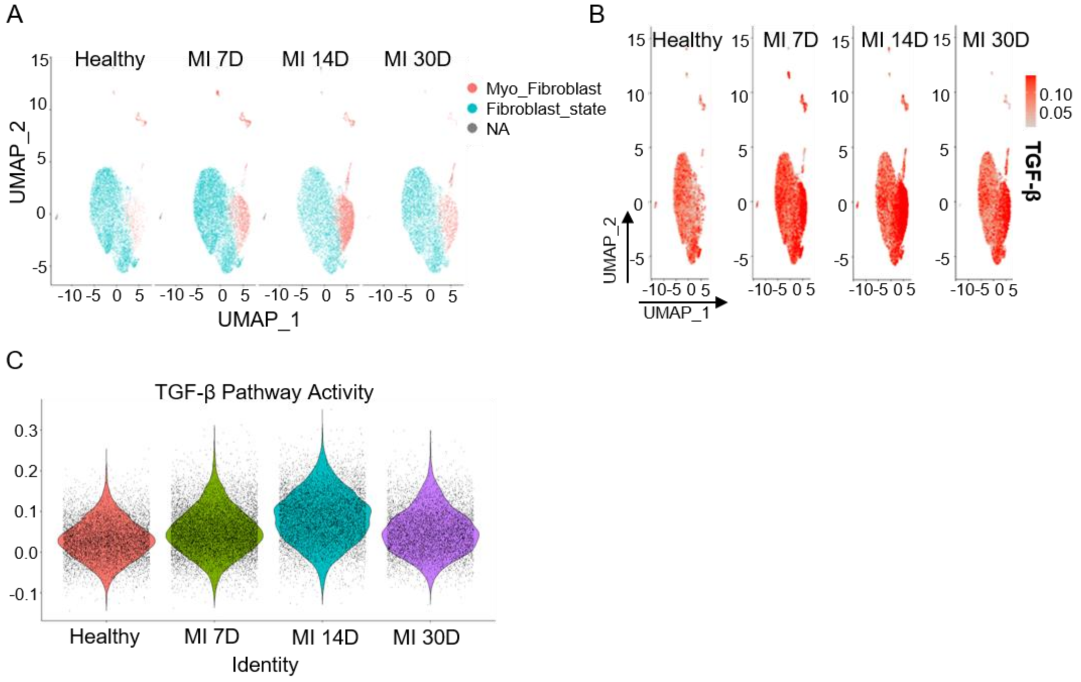


**Figure S2. Dynamic changes in key cell populations and key pathways.**

A, UMAP plot of fibroblast and myofibroblast across MI phases (0D, 7D, 14D and 30D).

B, Spatiotemporal expression of TGF-β pathway activity across four MI phases.

C, Violin plot of TGF-β pathway activity across MI phases.

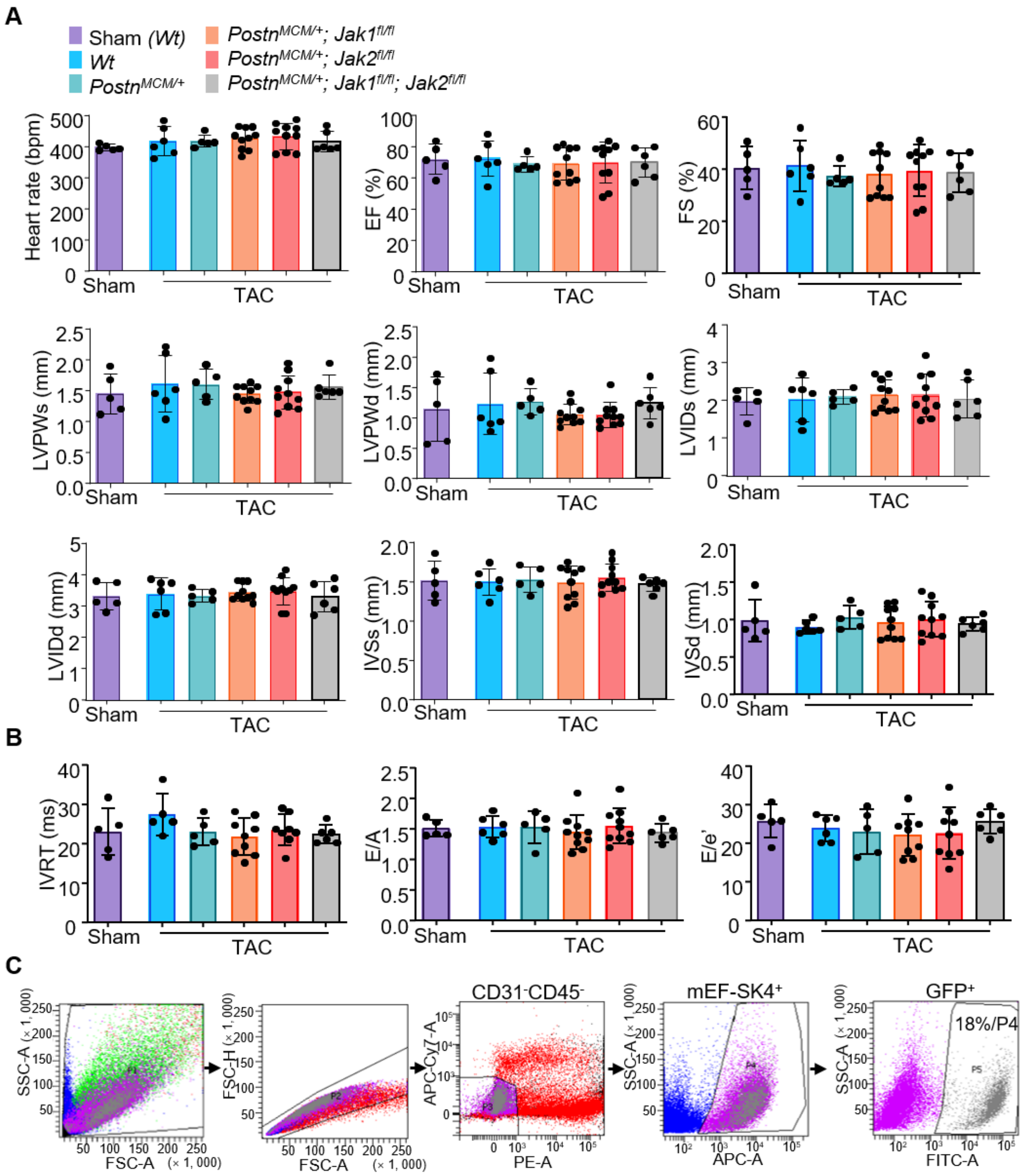


**Figure S3. Echocardiographic Assessment Reveals Preserved Cardiac Function in *Jak1/2* Knockout Mice Following Pressure Overload.**

A, Echocardiographic evaluation of cardiac systolic function and structural remodeling after TAC surgery. Left ventricular ejection fraction (LVEF %), fractional shortening (FS %), left ventricular internal diameter (LVID), left ventricular posterior wall (LVPW) and interventricular septal thickness (IVS) at end-systole and end-diastole were measured to assess functional and architectural changes.

B, Echocardiographic assessment of diastolic function following TAC. Key parameters included early (E) and late (A) diastolic mitral inflow velocities, E/A ratio, E peak to mitral annular tissue Doppler e' peak ratio (E/e' ratio) and isovolumic relaxation time (IVRT).

C, Flow cytometry gating scheme used to identify and characterize cardiac myofibroblast populations. One-way ANOVA was used followed by the Tukey multiple comparisons test. Individual data are presented as aligned dot plots, with the mean and SD.

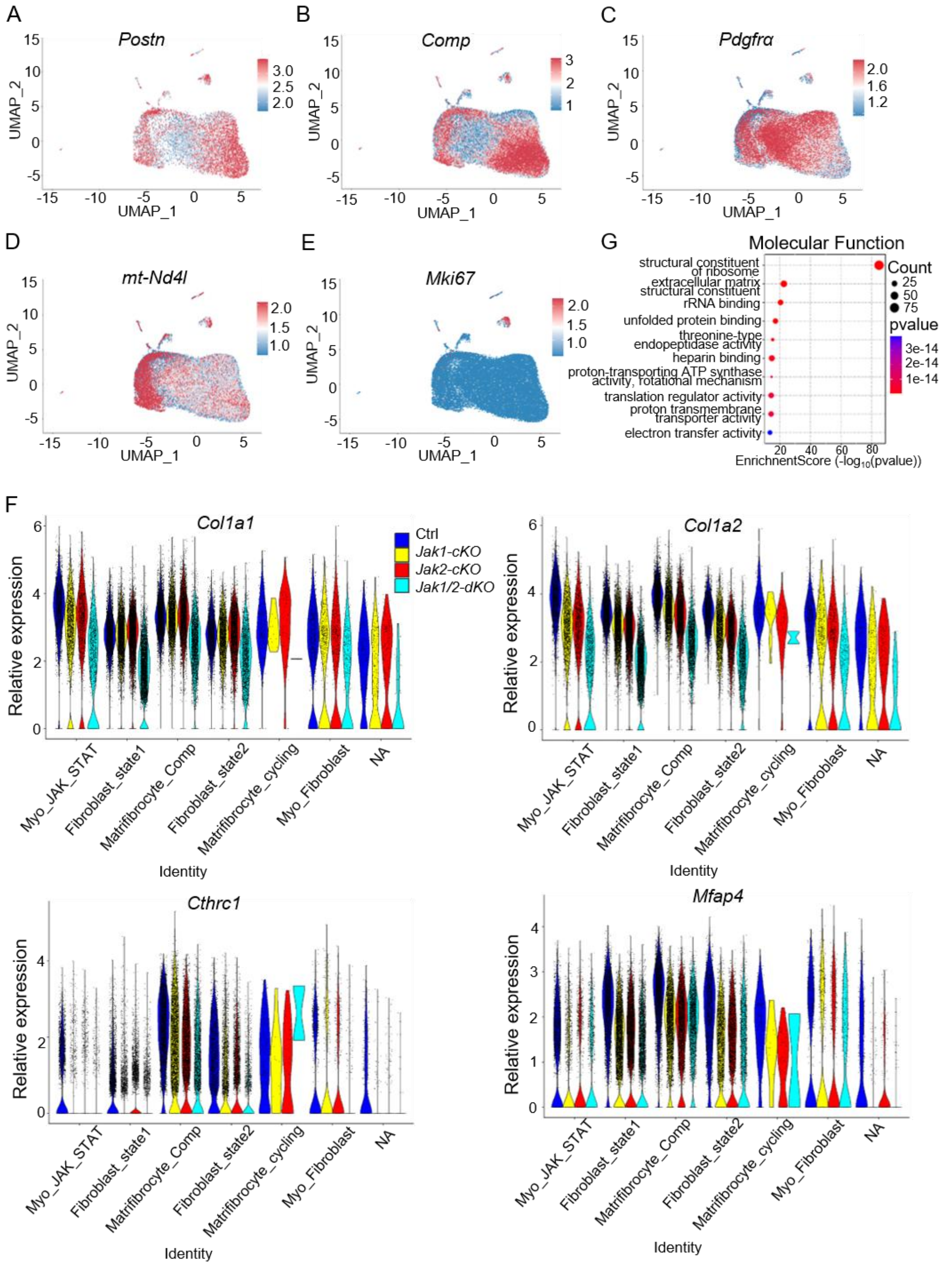


**Figure S4. Effects of *Jak1/2* Knockout on Expression of Fibrosis Core Genes.**

A through E, UMAP plot of key cell subpopulation marker genes.

F, Fibrosis-related genes (*Col1a1*, *Col1a2*, *Cthrc1* and *Mfap4*) expression in myofibroblasts by clusters as violin plot (y axis is normalized UMI levels).

G, Key pathways influencing Myo_JAK_STAT subpopulation differentiation.

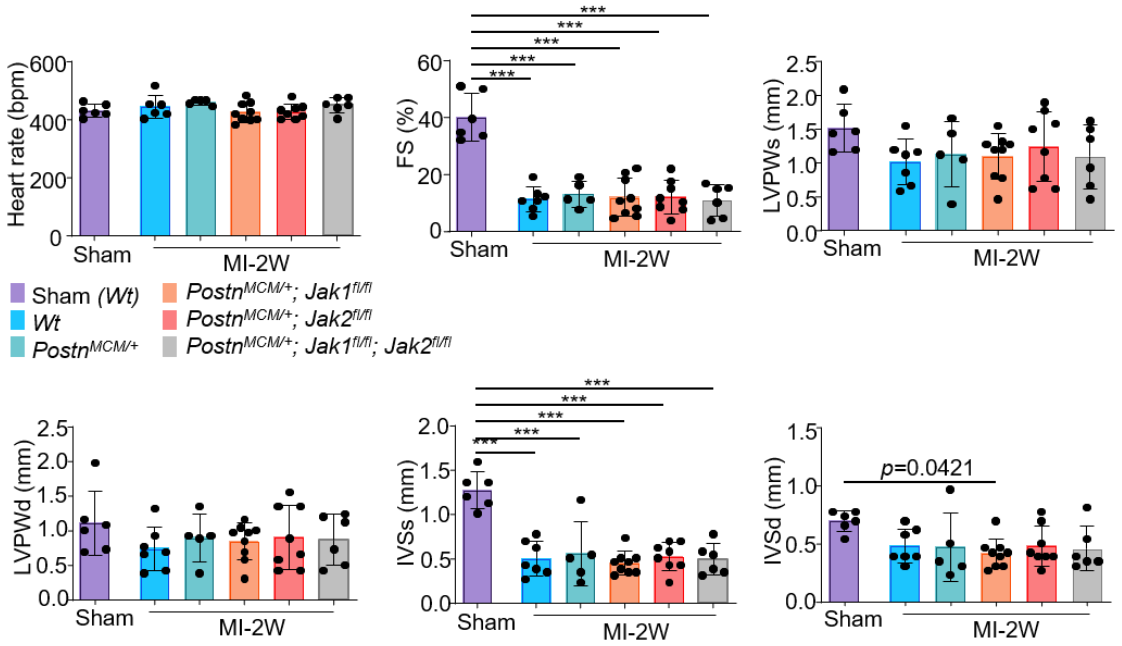


**Figure S5. Echocardiographic Assessment at 2 Weeks Post-MI Following Myofibroblast-Specific *Jak1/2* Knockout.**

Heart rate (HR), left ventricular fraction shortening (LVFS), left ventricular posterior wall in systole and diastole (LVPWs and LVPWd), interventricular septal thickness at end-systole and end-diastole (IVSs and IVSd) were quantified by echocardiography 14 days after MI in sham and knockout groups (n=6: 7: 5: 9: 8: 6). ***$P<0.001$ vs the sham group using 1-way ANOVA followed by Tukey multicomparisons test. Individual data are presented as aligned dot plots, with the mean and SD.

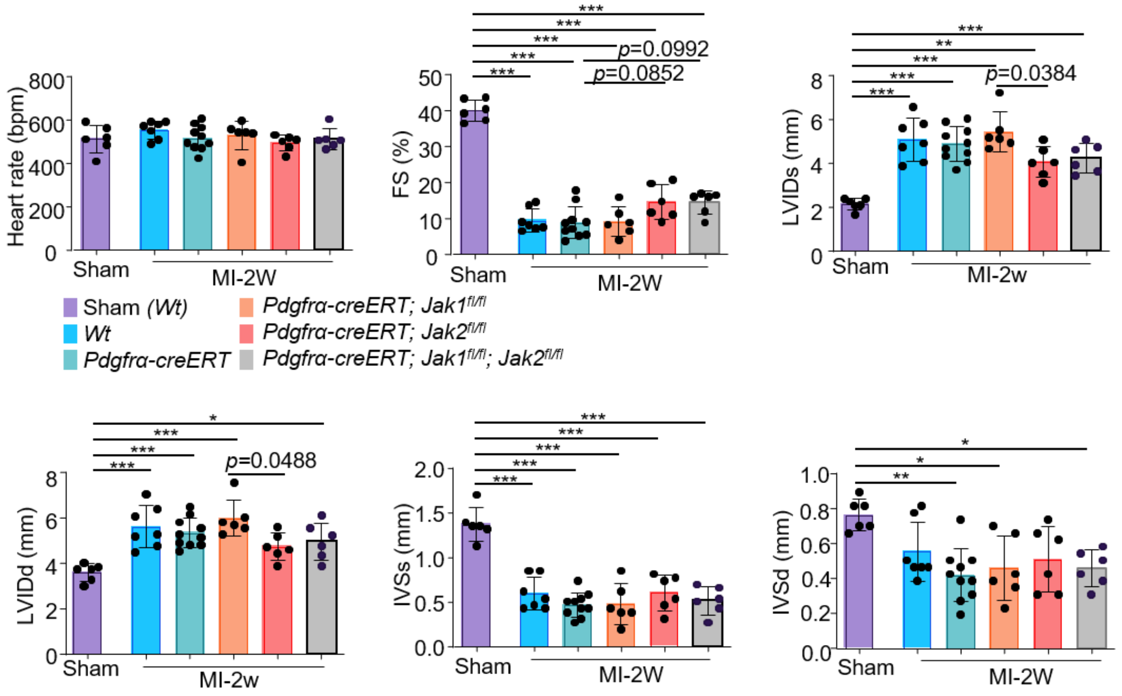


**Figure S6. Echocardiographic Assessment at 2 Weeks Post-MI Following Fibroblast-Specific *Jak1/2* Knockout.**

Heart rate (HR), left ventricular fraction shortening (LVFS), LV systolic and diastolic internal diameter (LVIDs and LVIDd), interventricular septal thickness at end-systole and end-diastole (IVSs and IVSd) were quantified by echocardiography 14 days after MI in sham and knockout groups (n=6: 7: 10: 6: 6: 6). **P*<0.05; ***P*<0.01; ****P*<0.001 vs the sham group using 1-way ANOVA followed by Tukey multicomparisons test. Individual data are presented as aligned dot plots, with the mean and SD.

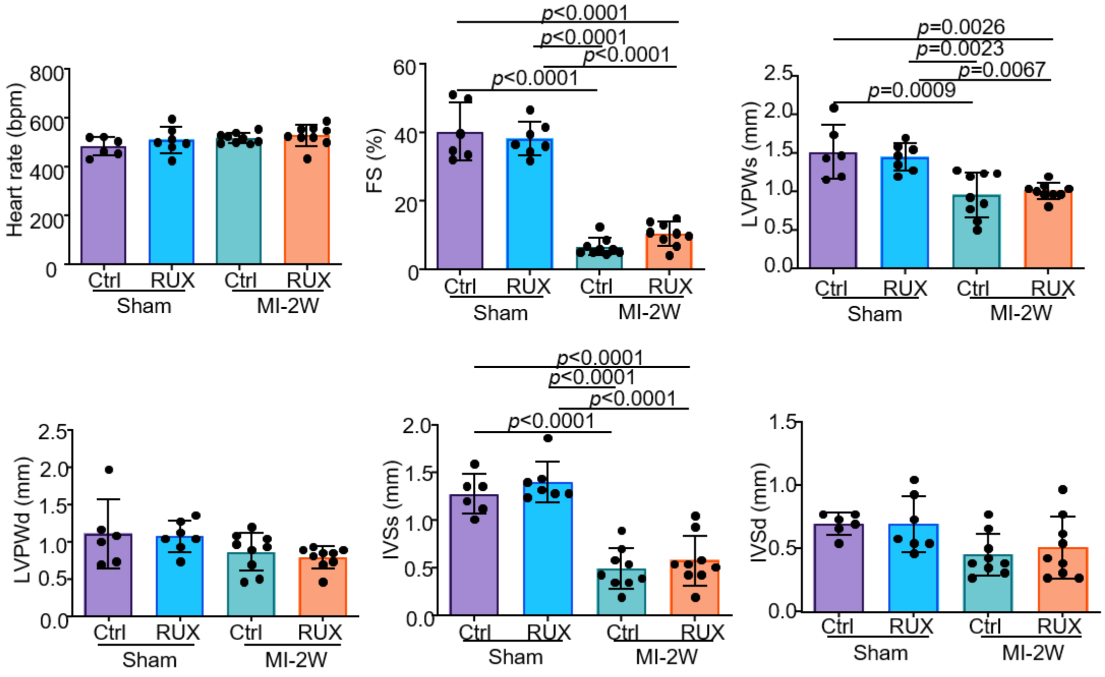


**Figure S7. Echocardiographic Assessment in Ruxolitinib-Treated Mice at 2 Weeks Post-Myocardial Infarction.**

Heart rate (HR), left ventricular fraction shortening (LVFS), left ventricular posterior wall in systole and diastole (LVPWs and LVPWd), interventricular septal thickness at end-systole and end-diastole (IVSs and IVSd) were quantified by echocardiography 14 days after MI in sham and knockout groups (n=6: 7: 9: 9).

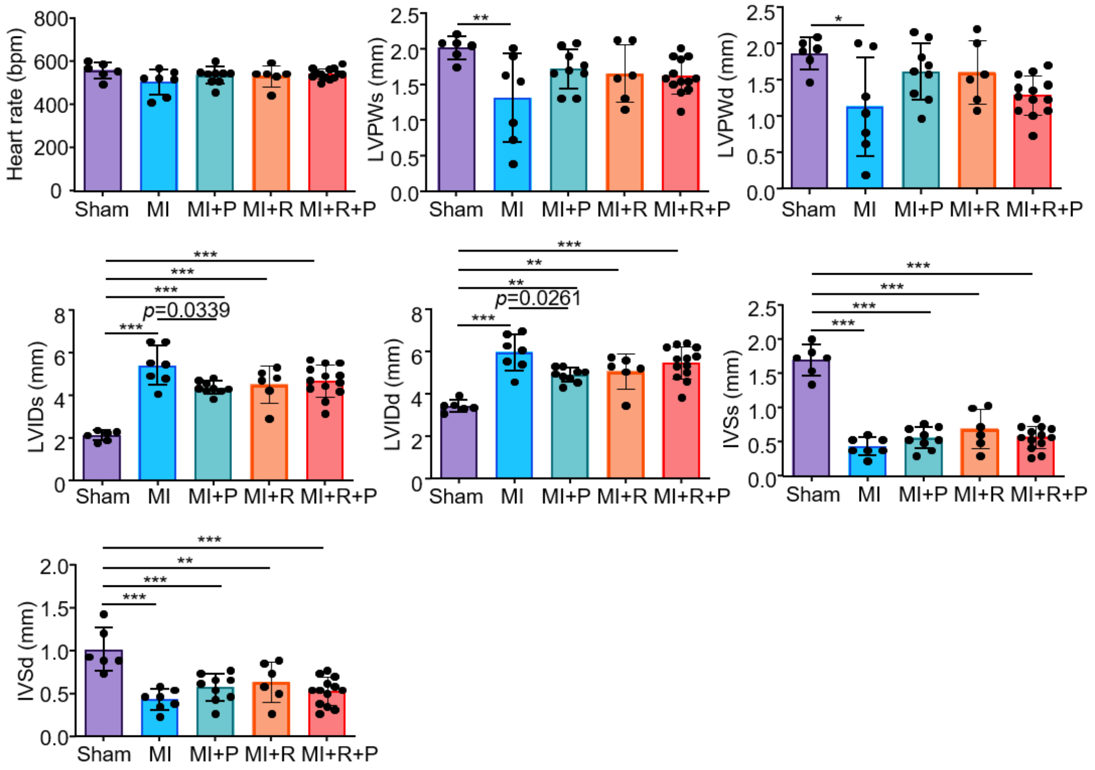


**Figure S8. Echocardiographic Assessment in Mice Post-Myocardial Infarction After Ruxilitnib and Pirfenidone Treatment.**

Heart rate (HR), left ventricular posterior wall in systole and diastole (LVPWs and LVPWd) and interventricular septal thickness at end-systole and end-diastole (IVSs and IVSd) were quantified by echocardiography 14 days after MI in sham and knockout groups (n=6: 7: 9: 6: 13). **P*<0.05; ***P*<0.01; ****P*<0.001 vs the sham group using 1-way ANOVA followed by Tukey multicomparisons test. Individual data are presented as aligned dot plots, with the mean and SD.
